# *C. elegans* interprets dietary quality through context-dependent serotonergic modulation

**DOI:** 10.1101/2025.01.05.631367

**Authors:** Likui Feng, Javier Marquina-Solis, Lishu Yue, Audrey Harnagel, Yarden Greenfeld, Cornelia I. Bargmann

**Affiliations:** Lulu and Anthony Wang Laboratory of Neural Circuits and Behavior, The Rockefeller University, New York, NY 10065, USA

## Abstract

Animals sense their metabolic needs to guide foraging decisions using neuronal pathways that are only partly understood. Here, we systematically investigate how foraging in the nematode *Caenorhabditis elegans* is influenced by its bacterial diet, *E. coli*. By screening *C. elegans* behavior on 3983 *E. coli* knockout strains, we identified 22 *E. coli* metabolic mutants that are aversive to *C. elegans* in a long-term foraging assay. These include the global metabolic regulator CRP and genes affecting cysteine synthesis, vitamin B6 synthesis, and iron uptake. Serotonin, a neurotransmitter associated with feeding in many animals, allows *C. elegans* to distinguish wild-type *E. coli* from these “mediocre” diets through bidirectional signaling. Serotonin produced by the ADF serotonergic neurons supports attraction to wild-type *E. coli* with the serotonin receptor genes *ser-4* and *ser*-*5*, whereas serotonin produced by the NSM serotonergic neurons differentially drives aversion to two mediocre diets through four serotonin receptor genes, *ser-1, ser-7, mod-1,* and *lgc-50.* Serotonin receptors act in multiple target neurons, including octopamine-producing neurons that suppress aversion across all diets. In addition, dopamine promotes aversion, in part by inhibiting octopaminergic neurons. These results reveal interactions between neuromodulatory circuits in the context-dependent evaluation of dietary quality.

## INTRODUCTION

Animals discriminate between high-quality and low-quality food to optimize their development, reproduction, and physiology. While food quality can be sensed through olfactory and gustatory cues, internally-sensed metabolic needs also drive food preferences. For example, an interoceptive dopaminergic circuit allows *Drosophila* to reject an incomplete diet lacking essential amino acids.^1^ Preferences can also change based on an animal’s physiology: newly eclosed female mosquitos feed on nectar, but after mating they prefer blood meals that support the growth of offspring.^2,3^ The full extent and nature of metabolic factors that influence food preference in animals are not known.^4,5,6^

The nematode *Caenorhabditis elegans* feeds on a variety of bacteria in its natural environment,^7^ but is typically cultivated on *E. coli* in the laboratory.^8^ Its nutritional requirements are well-characterized,^9,10^ and can be used to understand the influences of diet on its growth and physiology. For example, a metabolic pathway for alleviating vitamin B12 deficiency in *C. elegans* was discovered based on differential B12 availability in *E. coli* HT115(DE3) compared to the most commonly used *E. coli* food, OP50.^11–13^ This discovery inspired a number of studies of *C. elegans* growth on the 3983 *E. coli* strains in the Keio gene knockout library, identifying 244 *E. coli* genes required for rapid *C. elegans* development^14^ as well as *E. coli* genes that regulate *C. elegans* longevity,^15^ survival under mitochondrial stress,^16^ and resistance to the toxins 5-Fluoruracil, FUDR, camptothecin, and Aflatoxin B1.^17–19^

Its bacterial diet can also modulate *C. elegans* behavior. In addition to rapid chemotaxis to food-related metabolites and food ingestion through its pharynx,^20^ it has longer-term foraging behaviors that reflect its evaluation of food quality and quantity over minutes to hours.^21^ Over these timescales, animals display low-activity dwelling behaviors in high-quality food, and higher-activity roaming behaviors when food is limited.^22^ ^23–25^ At the extreme, animals will abandon a bacterial food source that is hard to eat, toxic, or pathogenic.^23,26–28^ Roaming, dwelling, and lawn-leaving (aversion) behaviors are under extensive neuromodulatory regulation by the biogenic amines serotonin,^25,29,30^ dopamine,^31,32^ tyramine,^33,34^ and octopamine,^35^ the neuroendocrine cytokine DAF-7,^36,37^ and the neuropeptides PDF-1 and FLP-1,^25,26^ ^38^ suggesting that long-term behaviors are directed by slow-acting neuromodulatory systems.

Serotonin links physiology and behavior in many animals. In mammals, serotonin is synthesized by neurons in the brainstem and by enteric neurons and enteroendocrine cells in the digestive system; it regulates locomotion, anxiety, and food responses including appetite, satiety, obesity, gut motility, and nausea.^39,40^ Similarly, the serotonergic system in *C. elegans* is implicated in a variety of food-related behaviors. The primary sources of serotonin are three pairs of neurons called NSM, ADF, and HSN. The NSM enteric neurons reside in the pharynx, detect food through proprioceptive endings, and release serotonin to stimulate acute feeding, locomotor slowing, and long-lasting dwelling behaviors.^41^ The ADF sensory neurons detect bacterial metabolites and regulate both acute feeding behaviors and learned avoidance of bacterial pathogens.^42,43^ The HSN neurons reside in the midbody and regulate egg-laying, locomotion, and dwelling behaviors.^44^ Serotonin acts on six receptors with different signaling mechanisms that are collectively expressed on nearly half of all *C. elegans* neurons; several receptors contribute to each serotonin-regulated behavior.^25,45^ The general organization of serotonin systems with a few serotonin-expressing neurons, multiple serotonin receptors, and widely distributed serotonin receptor-expressing neurons appears to be shared between mammals and *C. elegans.* As serotonin and other neuromodulators act extrasynaptically, these distributed systems create a challenge for circuit mapping. Understanding information flow requires detailed identification of the functional connections between serotonin- and serotonin-receptor expressing neurons in specific behavioral contexts.^45–47^

Here, we use the Keio *E. coli* gene knockout library to systematically identify “mediocre” bacterial diets that elicit aversion behavior in *C. elegans*. Pathway enrichment of these *E. coli* mutants suggests that *C. elegans* monitors dietary quality using a few primary bacterial signals: global bacterial metabolism (*crp*), cysteine biosynthesis, vitamin B6 biosynthesis, ferric iron transport, and bacterial inner membrane integrity. Using *C. elegans* genetics and circuit mapping, we find that serotonin has a dual role in assessing dietary quality: it promotes aversion from a mediocre diet and suppresses aversion from a high-quality diet through context-dependent roles in serotonin-producing neurons, serotonin receptors, and target neurons. In addition, dopamine promotes aversion, whereas octopamine suppresses aversion in all dietary contexts. Serotonin and dopamine act in part by modulating octopaminergic and tyraminergic cells, but serotonin also acts on other neuronal targets to regulate behavior. Our results reveal a distributed neuromodulatory network of biogenic amines, receptors, and target neurons that interpret dietary quality through cooperative and antagonistic effects on shared behaviors.

## RESULTS

### A genome-wide screen for *E. coli* mutants that elicit aversion behavior in *C. elegans*

We evaluated *C. elegans* preferences for various bacterial diets using a quantitative assay developed to characterize responses to toxic or pathogenic bacteria. 15-20 *C. elegans* L4 larvae are placed on a plate seeded with a small patch of bacterial food, and the location of the animals is tracked by video recording for 20 hours under constant environmental conditions (**Figure 1A**). Over 95% of animals remain within a patch of the standard bacterial food, *E. coli* OP50, throughout the 20-hour assay, whereas fewer than 10% of animals remain on patches of pathogenic *Pseudomonas aeruginosa* bacteria after 16-20 hours (**Figure 1B**). We defined an aversion ratio as the steady-state fraction of animals off the food patch 16-20 hours after the beginning of the assay (**Figure 1C**). Over 80% of animals remained on several wild-type *E. coli* strains throughout the 20-hour assay (**Figure 1B**).

**Figure 1.**
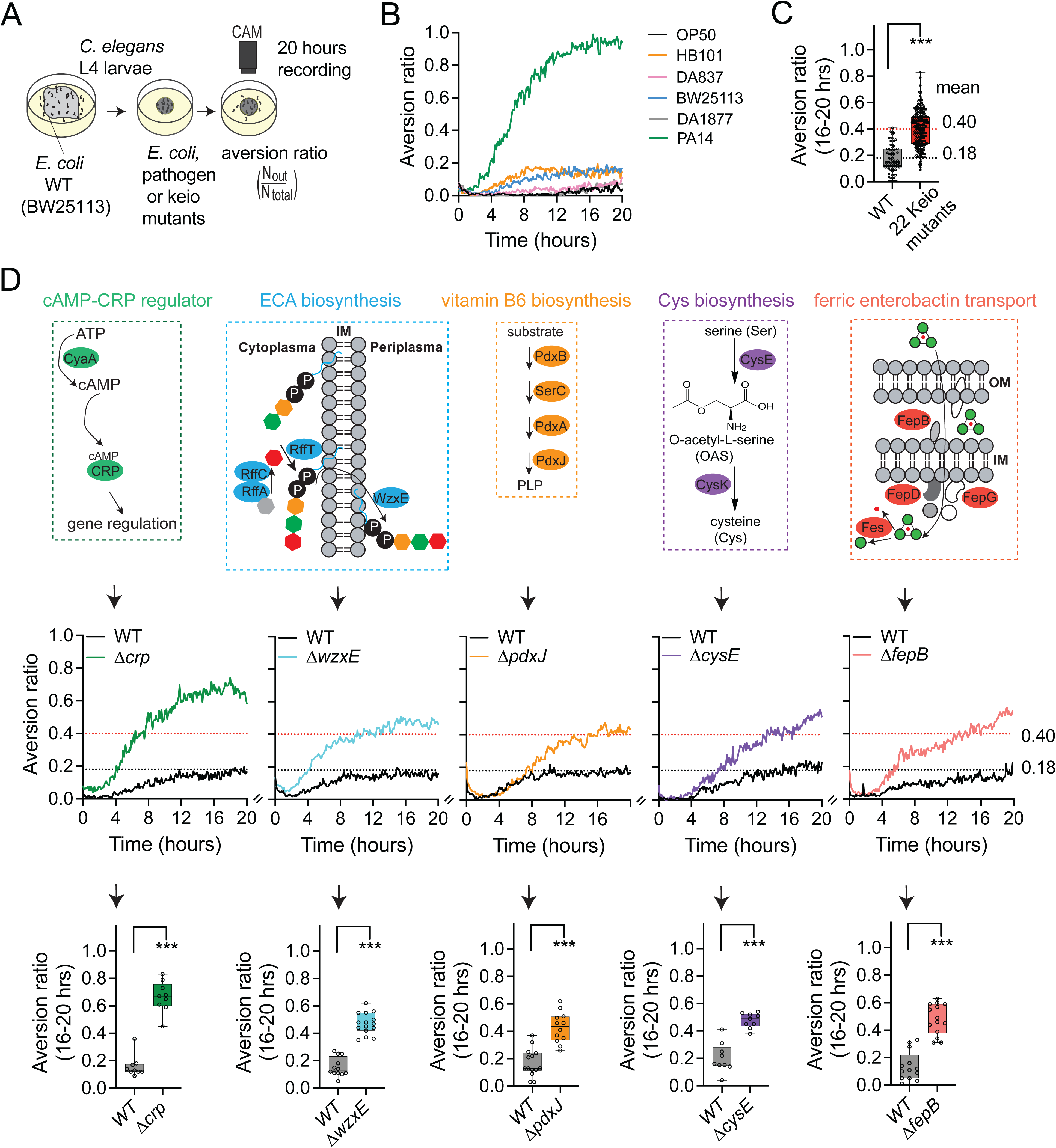
Genome-wide screen for *E. coli* mutants that induce aversion behavior in *C. elegans*. (A) Schematic depiction of aversion behavior screen. (B) Behavioral responses of wild-type *C. elegans* to different *E. coli* strains and to the pathogenic bacteria *Pseudomonas aeruginosa* PA14. Each trace represents the average of at least 8 individual assays per strain across 20 hours. (C) Comparison of pooled behavioral responses of wild-type *C. elegans* on WT BW25113 and the 22 Keio mutants identified from the whole-genome screen. The mean values for each group are shown. Black dashed line indicates mean value from WT group and red dashed line indicates mean value from pooled positive hints. (D) Simplified schematic depiction of identified metabolic pathways in *E. coli,* full aversion assays, and aversion ratios at 16-20 hours. Complete pathways are shown in Figure S1. Genes identified in this study are highlighted in color. cAMP, cyclic AMP. CRP, cAMP Receptor Protein. ECA, Enterobacterial Common Antigen. PLP, Pyridoxal 5’-phosphate. Ent, Enterobactin. IM, Inner Membrane. OM, Outer Membrane. Panels in the middle show aversion traces for one typical mutant chosen from each metabolic pathway. Panels at the bottom show quantification of averaged aversion ratios at 16-20 hours. Each data point indicates individual assay. ***<0.001, by one-way ANOVA, with Dunnett’s multiple correction.

To perform a systematic investigation of bacterial effects on *C. elegans* aversion behavior, we used the food patch assay to screen the *E. coli* Keio knockout collection^48^ of 3983 deletion mutants in the BW25113 genetic background. Following a prescreen that yielded 262 potentially aversive strains, we used quantitative video recording to identify 22 *E. coli* mutants that elicited significant aversion behavior (**Table 1**), with an average aversion ratio of 0.4 compared to 0.18 for the parental BW25113 strain (henceforth, wild-type) (**Figure 1C**). These 22 genes included multiple hits in five metabolic pathways: regulation of global transcription, biosynthesis of the enterobacterial common antigen (ECA), biosynthesis of the essential nutrients vitamin B6 and cysteine, and ferric iron transport (**Figure 1D and S1)**. These enriched pathways are described briefly below.

**Table 1.** *E. coli* mutants identified in the *C. elegans* behavioral aversion screen.

| Diet | Gene | Functions |
| --- | --- | --- |
| wild-type | WT (BW25113) | wild-type *E. coli*, K-12 strain |
| mediocre | *crp*, *cyaA* | cAMP-dependent global carbon response; *crp* encodes a DNA-binding transcriptional regulator |
|  | *wzxE*, *rffT*, *rffA*, *rffC* | enterobacterial common antigen (ECA) biosynthesis; *wzxE* encodes a lipid III ECA flippase |
|  | *pdxJ*, *pdxB*, *pdxA*, *serC* | pyridoxine (PN) and pyridoxine 5'-phosphate (PLP) biosynthesis; *pdxJ* encodes a PLP synthase |
|  | *cysE*, *cysK* | O-acetyl-L-serine (OAS) and cysteine (Cys) biosynthesis; *cysE* encodes a serine acetyltransferase |
|  | *fepB*, *fepD*, *fes*, *fepG* | ferric-enterobactin transport for iron bioavailability; *fepB* encodes an ABC transporter periplasmic binding protein |
|  | *fabH* | encodes a 3-oxoacyl-[acyl carrier protein] synthase in fatty acid biosynthesis |
|  | *tatB* | encodes a twin arginine translocation component inside inner membrane |
|  | *ynhG* | encodes a periplasmic L,D-transpeptidase |
|  | *alaS* | encodes an alanine-tRNA ligase |
|  | *aceF* | encodes a pyruvate dehydrogenase E2 subunit |
|  | *dnaJ* | encodes a chaperone protein |

In response to reduced availability of a preferred carbon source such as glucose, the *E. coli* adenylate cyclase CyaA catalyzes the production of cyclic AMP (cAMP), which is recognized and bound by CRP, the cAMP receptor protein transcription factor (**Figure 1D**).^49–52^ CRP regulates global bacterial metabolism in glucose-poor conditions such as Nematode Growth Medium. Δ*crp* and Δ*cyaA* elicit significant aversion in *C. elegans*; no aversion was induced by mutants for the cAMP phosphodiesterase CpdA or the CRP transcriptional cofactor cytR **(Figure S1A)**.

Most *Enterobacteriaceae* species including *E. coli* express enterobacterial common antigen (ECA) on the outer membrane, where it regulates bacterial cell shape and stress responses. The ECA backbone consists of undecaprenyl pyrophosphate-linked trisaccharide repeats that can be further modified by cyclization or conjugation with peptidoglycan or lipopolysaccharide. Four ECA biosynthetic mutants, Δ*wzxE*, *ΔrffT*, *ΔrffA* and Δ*rffC,* elicited aversive responses in *C. elegans.* RffA, RffC and RffT synthesize the terminal sugar moiety Fuc4NAc and attach it to the lipid-phosphate-disaccharide carrier, while the ECA flippase WzxE flips the resulting ECA unit across the inner membrane prior to ECA maturation^53^ (**Figure 1D**). No aversion was induced by mutants in earlier or later steps of ECA synthesis, suggesting that accumulation of an inner membrane intermediate, rather than the loss of ECA, drives *C. elegans* aversion behavior (**Figure S1B**).

Vitamin B6 cofactors such as pyridoxal, pyridoxine, pyridoxamine, and their phosphorylated forms are synthesized in *E. coli* by four essential enzymes (PdxB, SerC, PdxA, and PdxJ) and two pairs of redundant enzymes, Epd/GapA and PdxH/PdxK. Mutants in each of the essential enzymes induce aversion behavior, linking aversion to the loss of vitamin B6 cofactors (**Figure 1D and S1C)**. Vitamin B6 is an essential nutrient for *C. elegans.*^9^

In *E. coli,* cysteine is synthesized from serine through the serine acetyltransferase CysE, which converts L-serine to O-acetyl-L-serine (OAS), and the O-acetylserine sulfhydrylase CysK. CysE and CysK form a complex that allosterically activates serine to cysteine conversion;^54^ a second O-acetylserine sulfhydrylase, CysM, does not form a complex with CysE and plays a smaller role (**Figure 1D and S1D)**. The aversion induced by Δ*cysE* and Δ*cysK* but not Δ*cysM* suggests that low levels of OAS or cysteine drive aversion. Cysteine is not an essential amino acid for *C. elegans* growth,^9^ but cysteine limitation reduces synthesis of the protective redox metabolite glutathione.^55,56^

Fifteen enzymes support *E. coli* ferric iron uptake and release through the small-molecule siderophore enterobactin. The four mutants that elicit aversion in *C. elegans,* Δ*fepB,* Δ*fepD,* Δ*fepG,* and Δ*fes* (**Figure 1D and S1E**) accumulate a ferric iron-enterobactin complex in the *E. coli* cytoplasm or periplasm that has been proposed to be toxic to *C. elegans* mitochondria.^14,16^ These mutants elicit production of reactive oxygen species (ROS), but other ROS-producing mutants do not elicit aversion **(Figure S1E and S1F)**.

Individual mutants that can drive *C. elegans* aversion include Δ*fabH* (fatty acid biosynthesis), Δ*tatB* (twin arginine translocation component, Δ*ynhG* (L,D-transpeptidase), Δ*alaS* (alanine-tRNA ligase), Δ*aceF* (pyruvate dehydrogenase E2 subunit), and Δ*dnaJ* (a chaperone protein) **(Figure S1G and Table 1**). In summary, several “mediocre” *E. coli* diets can induce aversion behavior in *C. elegans*.

The relatively small number of genes and pathways represented suggests that aversion is an active, relatively specific behavior; in particular, over 200 of the *E. coli* deletion mutants that result in slow *C. elegans* growth did not elicit aversion in this assay.^40^

### *E. coli* mutants that elicit *C. elegans* aversion induce stress response pathways

We selected one *E. coli* mutant from each of the five biological pathways (Δ*crp*, Δ*wzxE*, Δ*pdxJ*, Δ*cysE* and Δ*fepB*) to examine in more detail. To refine our understanding of the bacterial mutants, we examined their effects on *C. elegans* molecular reporters of physiological stress.^27^ Expression of a *daf-7::GFP* reporter in ASJ sensory neurons is induced by bacterial pathogens and by food that is physically difficult for *C. elegans* to ingest.^36,37^ This reporter gene was significantly induced by Δ*crp,* Δ*wzxE,* and Δ*fepB* bacteria, but not by Δ*cysE* or Δ*pdxJ* bacteria, (**Figure 2A**). An *hsp-6::GFP* reporter for mitochondrial stress was also significantly induced by Δ*crp* and Δ*fepB* bacteria^14^ (**Figure 2B**). A *gst-4::GFP* reporter for oxidative stress was induced by Δ*cysE* and Δ*wzxE* bacteria (**Figure 2C**), but not by Δ*crp,* Δ*fepB,* or Δ*pdxJ*. These results indicate that mediocre diets selectively activate *C. elegans* stress responses. Several other *C. elegans* stress reporters were also induced by one or more of the mediocre bacterial strains (**Figure S2A-S2E**).

**Figure 2.**
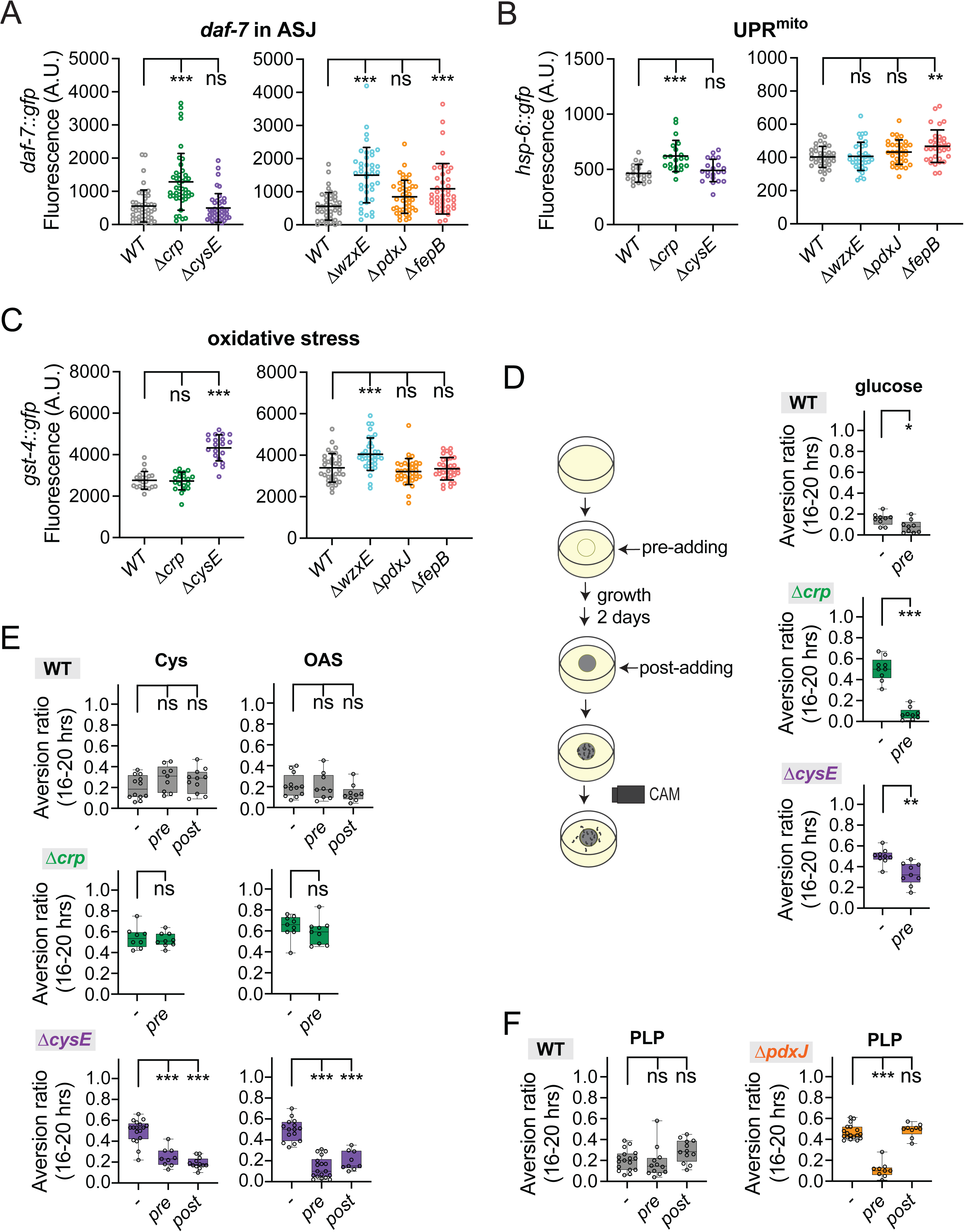
The mediocre diet-induced aversion in *C. elegans* is correlated with stress responses and metabolic states. (A-C) Effects of mediocre bacterial diets on *C. elegans* stress reporters. (B) Quantification of *daf-7::gfp* marker in ASJ sensory neurons. (C) Quantification of the mitochondrial UPR marker *hsp-6::gfp*. (D) Quantification of the oxidative stress marker *gst-4::gfp*. GFP fluorescence intensities quantified as Arbitrary Units (A.U.). Each data point indicates fluorescence from an individual animal. Error bars indicate mean ± SD. ns, not significant, **P<0.01, ***P<0.001 by One-Way ANOVA, corrected by Dunnett’s multiple comparisons. (D-F) Effects of chemical supplementation on aversion behavior. (D) Schematic diagram of chemical supplementation. Pre-adding indicates chemicals provided when bacteria grew on assay plates. Post-adding indicates chemicals supplied one hour before the behavioral assay. Pre-adding glucose strongly suppressed Δ*crp*-induced aversion. (E) Δ*cysE*-induced aversive responses were suppressed by pre- or post-addition of cysteine or the cysteine biosynthetic precursor OAS (F) Only pre-adding PLP suppressed Δ*pdxJ*-induced aversive responses. Final concentrations used for each chemical were glucose at 0.4%, Cys at 200 μM (pre-adding) or 50 μM (post-adding), OAS at 200 μM, and PLP at 40 μM. Each data point indicates individual assay. Results are shown with median ± quartiles in boxes and Min to Max whiskers. ns, not significant, *P<0.05, **P<0.01, ***P<0.001 by two-tailed, unpaired t test or One-Way ANOVA, corrected by Dunnett’s multiple comparisons.

We examined the effects of the selected *E. coli* mutants on *C. elegans* physiology as measured by developmental rate, brood size, feeding rate, colonization (a marker of bacterial pathogenesis), and lifespan **(Figure S2F)**. The strongest defects were observed on Δ*fepB*, which resulted in substantial developmental delay, reduced brood size, and colonization of *C. elegans;* these results align with the toxicity of this strain observed in previous screens.^14^ Δ*crp* resulted in slight developmental delay and reduced lifespan, and the other *E. coli* mutants caused only small alterations across one or more assays **(Figure S2F).**

The effects of the metabolic mutants on *C. elegans* behavior could be either direct readouts of the relevant bacterial pathways, or indirect effects of the mutations on other metabolic processes in the bacteria. To distinguish between these possibilities, we supplemented *E. coli* mutants with relevant metabolites and assessed aversion behavior. Addition of glucose during bacterial growth, which should bypass the metabolic function of CRP, fully suppressed *C. elegans* aversion to Δ*crp* mutants (**Figure 2D**). Addition of cysteine or its precursor OAS suppressed *C. elegans* aversion to Δ*cysE* either before or after the growth of the bacteria, suggesting that dietary cysteine deficiency caused aversion in *C. elegans* (**Figure 2E**). Vitamin B6 metabolites only suppressed *C. elegans* aversion when they were administered to the Δ*pdxJ* mutant or other mutants during bacterial growth, prior to the assay (**Figure 2F and S2G)**. These results suggest that aversion to Δ*pdxJ* may be an indirect consequence of a change in bacterial metabolism, not a direct effect of vitamin B6 deficiency. Similarly, vitamin B6 biosynthetic mutants confer *C. elegans* resistance to the toxin 5-fluorouracil by eliciting global changes in bacterial one-carbon and ribonucleotide metabolism that indirectly affect *C. elegans*.^17^

The results of these experiments suggest that at least two different bacterial qualities can drive *C. elegans* aversion: a metabolic change that induces mitochondrial stress (Δ*crp,* Δ*fepB*) and a metabolic change that induces oxidative stress (Δ*cysE);* both processes may contribute to aversion from Δ*wzxE*. While more remains to be discovered about *C. elegans* interactions with bacterial metabolism, Δ*crp* and Δ*cysE* were used as representative examples of aversive bacteria of two different classes.

### *C. elegans* evaluates opposing food qualities with distinct classes of serotonergic neurons

To ask how *C. elegans* distinguishes between bacteria in the aversion assay, we began with serotonin, a neuromodulator that regulates multiple food-related behaviors as well as avoidance to pathogenic bacteria.^25,27,42,57,58^ Animals mutant for the tryptophan hydroxylase gene *tph-1* lack all neuronally-synthesized serotonin.^59,60^ We found that *tph-1* mutants had reduced aversion to both Δ*crp* and Δ*cysE* mediocre diets, while retaining apparently normal responses to wild-type bacteria (**Figure 3A**).

**Figure 3.**
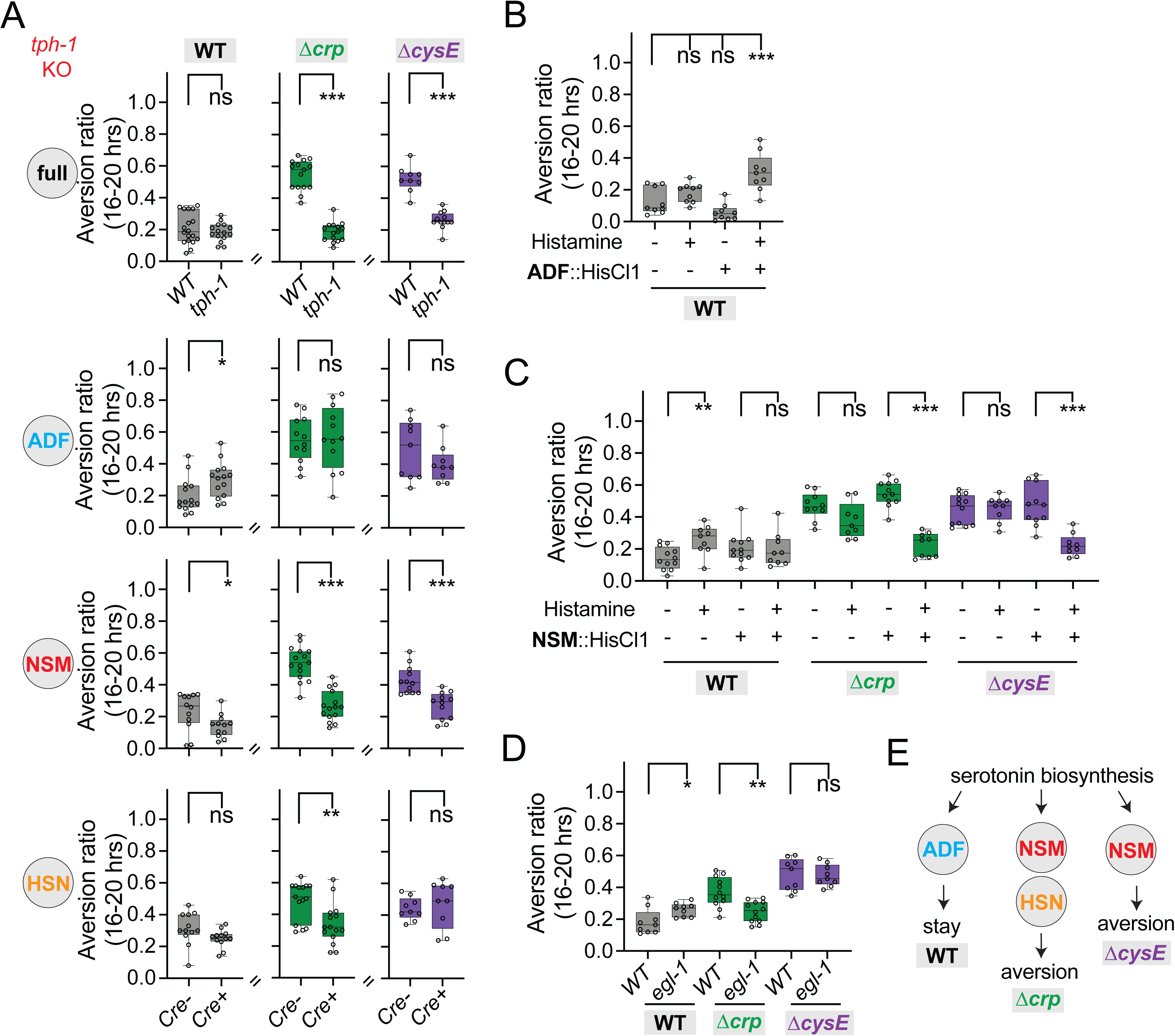
*C. elegans* evaluates opposing food qualities through different serotonergic neurons. (A) Aversion behaviors of *tph-1* null and neuron-specific knockout mutants on three diets. *tph-1* is expresses in three major serotonergic neuron classes ADF, NSM and HSN, and encodes a tryptophan hydroxylase that catalyzes the first step in serotonin biosynthesis. Cell-specific knockout of a single-copy genomic *tph-1* insertion was achieved by Cre-Lox recombination.^25^ (B) ADF silencing with a HisCl transgene and histamine induces aversion to wild-type *E. coli* BW25113. (C) NSM silencing with a HisCl transgene and histamine suppresses aversion to Δ*crp* and Δ*cysE* diets. (D) HSN-deficient *egl-1(gf)* mutants show decreased aversion to the Δ*crp* diet. (E) Schematic model of serotonergic neuron functions. For all panels, each data point indicates individual assay. Results are shown with median ± quartiles in boxes and Min to Max whiskers. ns, not significant, *P<0.05, **P<0.01, ***P<0.001 by two-tailed, unpaired t test (panels A, C and D) or One-Way ANOVA, corrected by Dunnett’s multiple comparisons (panel B).

Three classes of *tph-1-*expressing neurons synthesize serotonin in *C. elegans* hermaphrodites, ADF, NSM and HSN. To ask which neurons regulate aversion, we eliminated *tph-1* in individual neuron classes using Cre/lox recombination, ^25^ and tested the resulting animals on wild-type, Δ*crp* and Δ*cysE* diets. Remarkably, each of the three serotonin-producing neurons affected aversion behaviors (**Figure 3A**). Depletion of serotonin synthesis in ADF neurons resulted in a small but significant increase in aversion from wild-type bacteria, depletion of serotonin synthesis in NSM neurons decreased aversion from both Δ*crp* and Δ*cysE* mediocre diets, and depletion of serotonin synthesis in HSN neurons decreased aversion from the Δ*crp* diet. These results suggest that serotonin can either increase or decrease aversion behavior, depending on the neuronal serotonin source and the dietary context.

To validate these results with an orthogonal approach, we manipulated individual serotonergic neurons and tested the resulting strains on the three bacterial diets. Acute chemogenetic silencing of ADF neurons using a histamine-gated chloride channel (HisCl1) transgene^61^ and exogenous histamine significantly increased aversion from wild-type bacteria (**Figure 3B**), resembling the *tph-1* knockout in ADF. Acute chemogenetic silencing of NSM neurons suppressed aversion to the mediocre Δ*crp* and Δ*cysE* diets, recapitulating the effects of the *tph-1* knockout in NSM (**Figure 3C**). Finally, a strain in which the HSN neurons were genetically ablated (*egl-1(gf))* had diminished aversion to the mediocre Δ*crp* diet, resembling animals in which HSN did not synthesize serotonin (**Figure 3D**). These results confirm that serotonergic ADF neurons prevent aversion from wild-type diets, whereas serotonergic NSM and to a lesser extent HSN neurons promote aversion from mediocre Δ*crp* and Δ*cysE* diets (**Figure 3E**).

### The serotonin receptor SER-5 suppresses aversion from a wild-type diet together with the biogenic amine octopamine

*C. elegans* has six known serotonin receptors: two serotonin-gated ion channels (MOD-1, a chloride channel, and LGC-50, a cation channel), and four G-protein coupled receptors (SER-5 (Gs-coupled), SER-7 (Gs-coupled), SER-1 (Gq-coupled) and SER-4 (Gi/o-coupled)) (**Figure 4A**).^59^ Collectively, these receptors are expressed in almost half of all neuronal classes, and typically act cooperatively in foraging and feeding behaviors.^45^ We tested mutants for each serotonin receptor for aversion on the wild-type diet as well as mediocre Δ*crp* and Δ*cysE* diets; for clarity, the results are discussed separately on each diet below.

**Figure 4.**
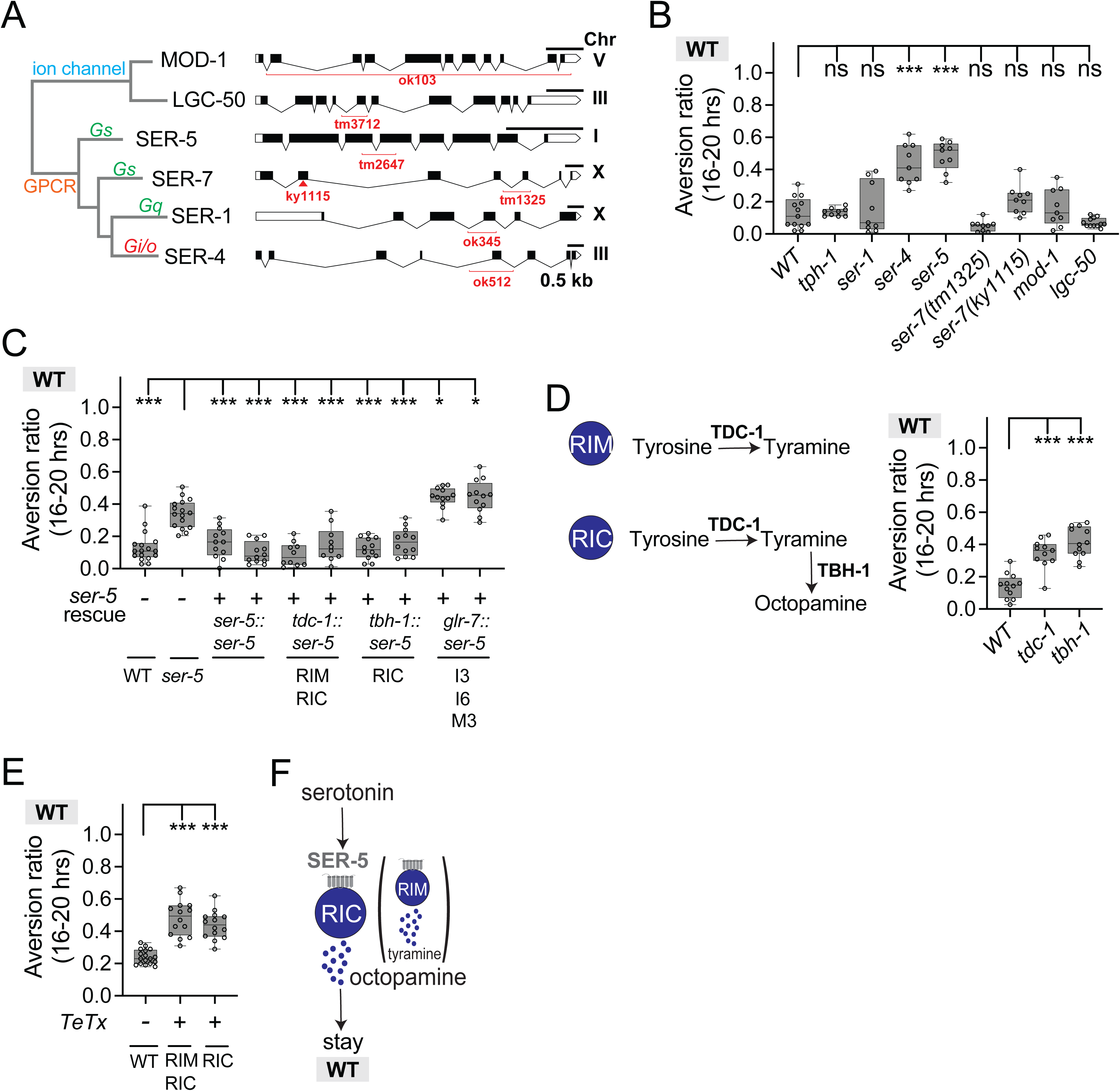
The serotonin receptor SER-5 suppresses aversion from a wild-type diet. (A) Schematic diagram of six serotonin receptors in *C. elegans,* with relevant G-protein for GPCRs indicated. Alleles for each serotonin receptor mutants are highlighted in red on genomic map. (B) Behavioral responses of wild-type, *tph-1*, and serotonin receptor mutant animals on the wild-type BW25113 diet. *ser-4* and *ser-5* mutants showed significant aversion. (C) Aversion behavior of *ser-5* mutants expressing *ser-5* cDNA from its endogenous promoter (*ser-5*) or promoters for RIM and RIC (*tdc-1*), RIC (*tbh-1*), or pharyngeal neurons (*glr-7*). Two transgenic lines were tested for each rescue plasmid. (D) Aversion behavior of *tdc-1* and *tbh-1* mutants on BW25113. TDC-1 (a tyrosine decarboxylase) in both RIM and RIC neurons catalyzes tyrosine to produce tyramine (TA), which is converted by TBH-1 (a tyramine b-hydroxylase) to form octopamine (OA) in RIC neurons. (E) Expression of the tetanus toxin light chain in RIM and/or RIC to block tyramine and octopamine synaptic release recapitulated phenotypes of *tdc-1* and *tbh-1* mutants on BW25113. (F) Schematic model of SER-5 function. For (B) to (E), each data point indicates individual assay. Results are shown with median ± quartiles in boxes and Min to Max whiskers. ns, not significant, *P<0.05, ***P<0.001 by One-Way ANOVA, corrected by Dunnett’s multiple comparisons.

Two of the six serotonin receptor mutants, *ser-4* and *ser-5,* showed strikingly enhanced aversion to the wild-type diet, while the mutants in the other four receptor genes were normal (**Figure 4B**). Previous studies suggested that serotonin release from ADF neurons activates the excitatory SER-5 receptor in a variety of neurons to regulate sensory responses, food intake, and longevity.^62–64^ We found that a *ser-5* cDNA rescued behavior on the wild-type diet when expressed from its endogenous promoter or from promoters for *tdc-1* (expressed in RIM and RIC neurons) or *tbh-1* (expressed in RIC neurons) (**Figure 4C**). Expression in pharyngeal neurons implicated in feeding did not rescue behavior on the wild-type diet (**Figure 4C**).

The RIM and RIC neurons express tyrosine decarboxylase *(tdc-1)* to produce the neurotransmitter tyramine, and RIC also expresses tyramine beta-hydroxylase *(tbh-1)* that converts tyramine into octopamine. *tdc-1* mutants lack both tyramine and octopamine, whereas *tbh-1* mutants lack octopamine and accumulate excess tyramine.^33^ We examined the behaviors of *tdc-1* and *tbh-1* mutant animals on wild-type diets. Like *ser-5* mutant animals, both *tdc-1* and *tbh-1* mutants had substantial aversion from the wild-type diet (**Figure 4D**). Expressing the tetanus toxin light chain to block synaptic release from RIC neurons or from RIM and RIC neurons also induced aversion (**Figure 4E**).

Together, these results suggest that serotonin from ADF acts on SER-5 to enhance the activity of tyramine and octopamine-producing cells that suppress aversion from a high-quality diet (**Figure 4F**). The combination of strong *ser-5* rescue in RIC alone (**Figure 4C**) and equivalent levels of aversion induced by *tbh-1* and *tdc-1* mutations (**Figure 4D**) suggest that octopamine from RIC neurons is the primary signal that suppresses aversion from wild-type bacteria (**Figure 4F**) but tyramine may also have a role (see below).

### The serotonin receptor SER-7 and dopamine promote aversion from a mediocre Δ*crp* diet

The serotonin receptor mutants *ser-7, mod-1,* and *lgc-50* had reduced aversion from the mediocre Δ*crp* diet (**Figure 5A**). Previous studies have shown that the excitatory SER-7 receptor acts in enteric neurons of the pharynx to regulate subtle aspects of feeding.^24^ We found that a *ser-7* cDNA rescued aversion on the mediocre Δ*crp* diet when expressed from its endogenous promoter or a *flp-21* promoter, which is expressed in overlapping pharyngeal neurons (**Figure 5B**).^24^ Among these neurons, *ser-7* expression in the I1 pharyngeal neuron partly rescued aversion from the mediocre Δ*crp* diet, but expression in several other neurons did not (**Figure 5B** and **5C**). These results suggest that the I1 pharyngeal neuron, likely responding to serotonin released from the NSM pharyngeal neurons, is part of an enteric circuit that promotes aversion from the mediocre Δ*crp* diet (**Figure 5D**). Additional *ser-7, mod-1,* and *lgc-50*-expressing neurons that affect aversion from Δ*crp* remain to be identified.

**Figure 5.**
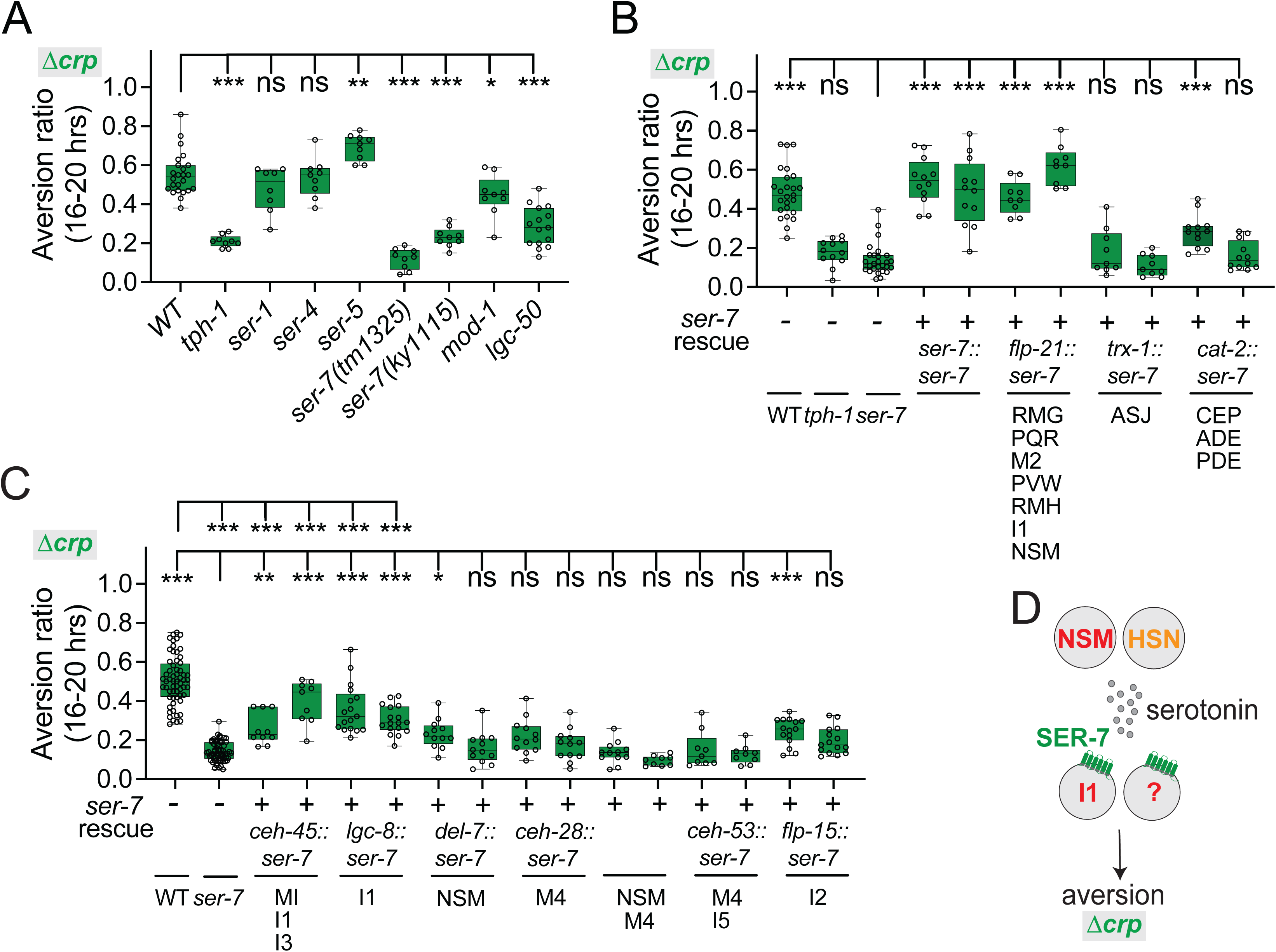
The serotonin receptor SER-7 drives aversion from the mediocre Δ*crp* diet. (A) Behavioral responses of wild-type, *tph-1*, and serotonin receptor mutant animals on the mediocre Δ*crp* diet. Aversion is diminished in two *ser-7* null mutants and in *mod-1* and *lgc-50* mutants. (B) Aversion behavior of *ser-7* mutants expressing *ser-7* cDNA from its endogenous promoter (*ser-7)*, or promoters for pharyngeal neurons (*flp-21*), ASJ (*trx-1*), or dopaminergic neurons (*cat-2*). Two transgenic lines were tested for each plasmid. (C) Aversion behavior of *ser-7* mutants with *ser-7* rescue in the I1 neuron and other sets of pharyngeal neurons. Two transgenic lines were tested for each rescue plasmid. (D) Schematic model of SER-7 serotonin signaling in aversion from the mediocre Δ*crp* diet. For (A), (B), and (C), each data point indicates individual assay. Results are shown with median ± quartiles in boxes and Min to Max whiskers. ns, not significant, *P<0.05, **P<0.01, ***P<0.001 by One-Way ANOVA, corrected by Dunnett’s (panel A and B), or Tukey’s (panel C) multiple comparisons.

Additional insight into aversion from the Δ*crp* diet came from examination of the neurotransmitter dopamine. Animals mutant for the dopamine biosynthetic enzyme *cat-2* (tyrosine-3-monooxygenase) had diminished aversion to the Δ*crp* mediocre diet (**Figure 6A**). *cat-2* is expressed in three classes of ciliated sensory neurons, CEP, ADE, and PDE. Expression of a *cat-2* cDNA from its own promoter or a promoter that was selectively expressed in CEP sensory neurons rescued aversion on the Δ*crp* diet (**Figure 6A**).

**Figure 6.**
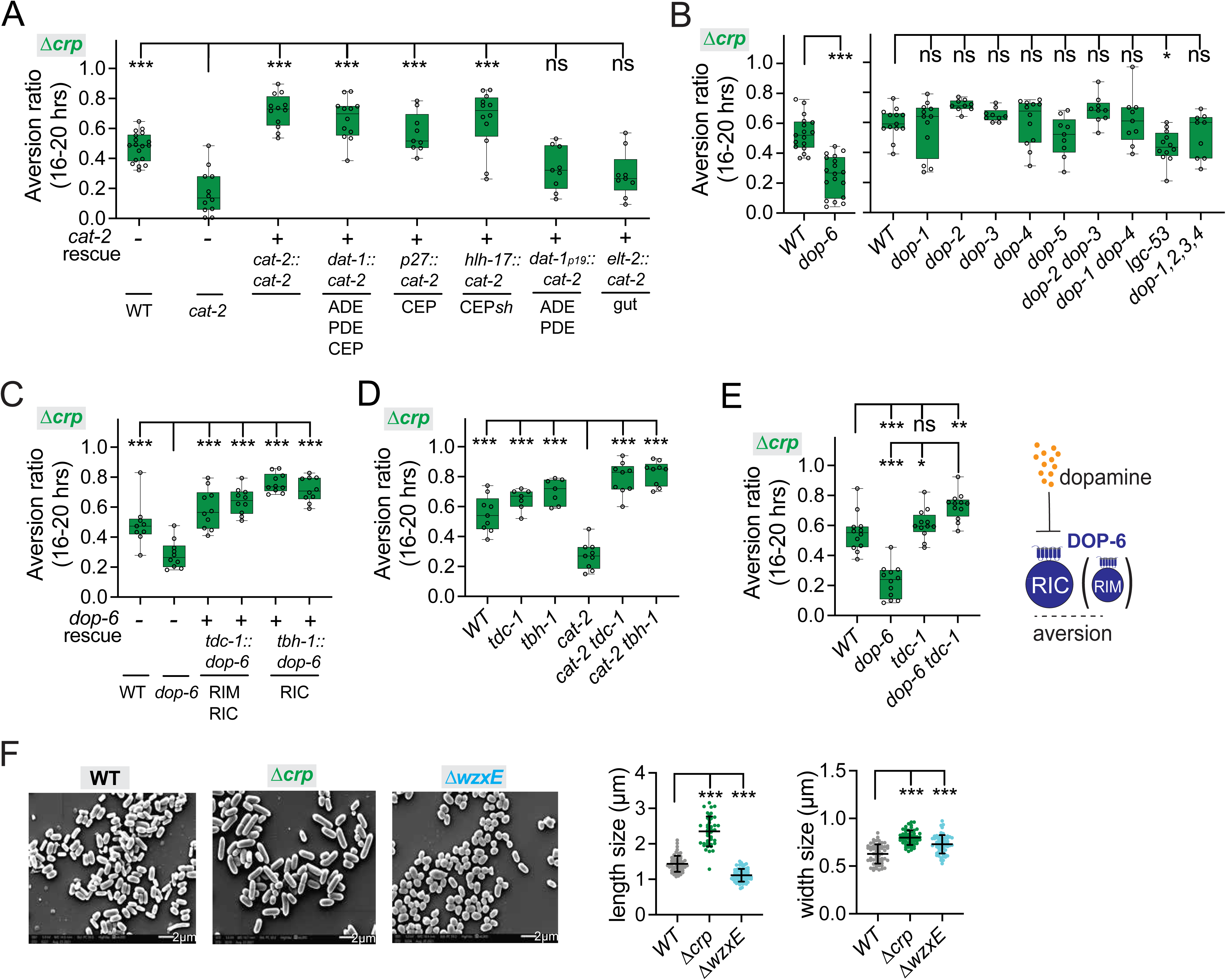
Aversion from the mediocre Δ*crp* diet requires dopamine. (A) Aversion behavior of *cat-2* mutants expressing *cat-2* genomic DNA from its the endogenous promoter (*cat-2*), or promoters for all dopaminergic neurons (*dat-1)*, CEP neurons (*p27*), CEP glial cells (*hlh-17*), ADE and PDE neurons (*dat-1p19*), or intestine (*elt-2*) on the mediocre *Δcrp* diet. (B) Aversion behavior of WT and dopamine receptor mutants on the mediocre *Δcrp* diet. (C) Aversion behavior of *dop-6* mutants with *dop-6* rescue in *tdc-1-* or *tbh-1-* expressing cells on the mediocre *Δcrp* diet. (D) Genetic interaction of *cat-2* with *tdc-1* or *tbh-1* (tyramine and/or octopamine biosynthesis mutants) on the mediocre *Δcrp* diet. *tdc-1* or *tbh-1* mutations restored behavioral aversion in the *cat-2* mutant background. (E) Genetic interaction of *dop-6* with *tdc-1* on the mediocre *Δcrp* diet, and schematic model of dopamine signaling. Stopped and dotted lines represent inhibition of RIC and octopamine. (F) Scanning electron micrographs of wild-type BW25113 bacteria, *Δcrp* bacteria, and *ΔwxzE* bacteria (left) and quantification of bacterial length and width (right, with SEM). Scale bar is 2 μm. For (A) to (E), each data point indicates individual assay. Results are shown with median ± quartiles in boxes and Min to Max whiskers. ns, not significant, *P<0.05, **P<0.01, ***P<0.001 by One-Way ANOVA, corrected by Dunnett’s multiple comparisons (panel A, B and C), two-tailed, unpaired t test (panel B, *dop-6)* or Tukey’s multiple comparisons (panel D and E). For (F), each data point indicates individual bacterial cell. ***P<0.001 by two-tailed, Mann-Whitney test.

Turning to dopamine receptors, animals mutant for the inhibitory (Gi/o-coupled) dopamine receptor DOP-6, but not other dopamine receptors, had reduced aversion to the Δ*crp* mediocre diet, like *cat-2* mutants (**Figure 6B**). Previous studies have shown that DOP-6 can inhibit the activity of RIC neurons.^64^ We found that expression of a *dop-6* cDNA in the RIC neuron, or in RIM and RIC neurons, rescued aversion from the mediocre Δ*crp* diet (**Figure 6C**).

To further explore the dopamine circuit, we used double mutants to ask how dopamine interacts with RIM and RIC neurotransmitters. Animals mutant for the RIM/RIC neurotransmitter enzymes *tdc-1* or *tbh-1* avoided the mediocre Δ*crp* diet (**Figure 6D**), as did *cat-2 tdc-1* and *cat-2 tbh-1* double mutants, unlike *cat-2* single mutants (**Figure 6D**). *tdc-1 dop-6* double mutants also avoided the mediocre Δ*crp* diet, as expected if DOP-6 inhibits tyramine and octopamine function from RIM and RIC (**Figure 6E**). As RIC expression was sufficient to rescue *dop-6,* and *cat-2* aversion defects were fully suppressed by *tbh-1,* it is likely that octopamine from RIC neurons can suppress aversion from the Δ*crp* diet, but is inhibited by dopamine.

The CEP neurons that produce dopamine are implicated in mechanosensory detection of bacteria.^65^ To ask whether Δ*crp* bacteria might have unusual mechanosensory features, we examined these bacteria by scanning electron microscopy. Under our growth conditions, Δ*crp* cells were significantly larger than wild-type *E. coli* (**Figure 6F**). A second bacterial strain with morphological alterations was Δ*wzxE,* the ECA biosynthesis mutant (**Figure 6F**). These defects may be related to the defective outer membrane structures of Δ*wzxE* mutants. Like aversion to the Δ*crp* diet, aversion from the Δ*wzxE* diet required *tph-1, cat-2,* and *ser-7* genes **(Figure S3).**

In summary, aversion from the mediocre Δ*crp* diet involves antagonistic interactions between serotonin and dopamine, which promote aversion, and octopamine, which suppresses aversion. Cell-specific rescue experiments identify NSM, HSN, CEP, I1, and RIC neurons as regulators of Δ*crp* aversion.

### Two serotonin receptors, SER-1 and MOD-1, promote aversion from a mediocre Δ*cysE* diet

The serotonin receptor mutants *ser-1* and *mod-1* had reduced aversion from the mediocre Δ*cysE* diet, like *tph-1* mutants (**Figure 7A** and **7B**). Previous studies showed that the excitatory SER-1 receptor regulates nociception through its action in RIC neurons.^66^ We found that a *ser-1* cDNA rescued aversion from the Δ*cysE* diet when expressed from its endogenous promoter or when expressed specifically in RIA neurons, but did not rescue when expressed in RIC neurons (**Figure 7C**).

**Figure 7.**
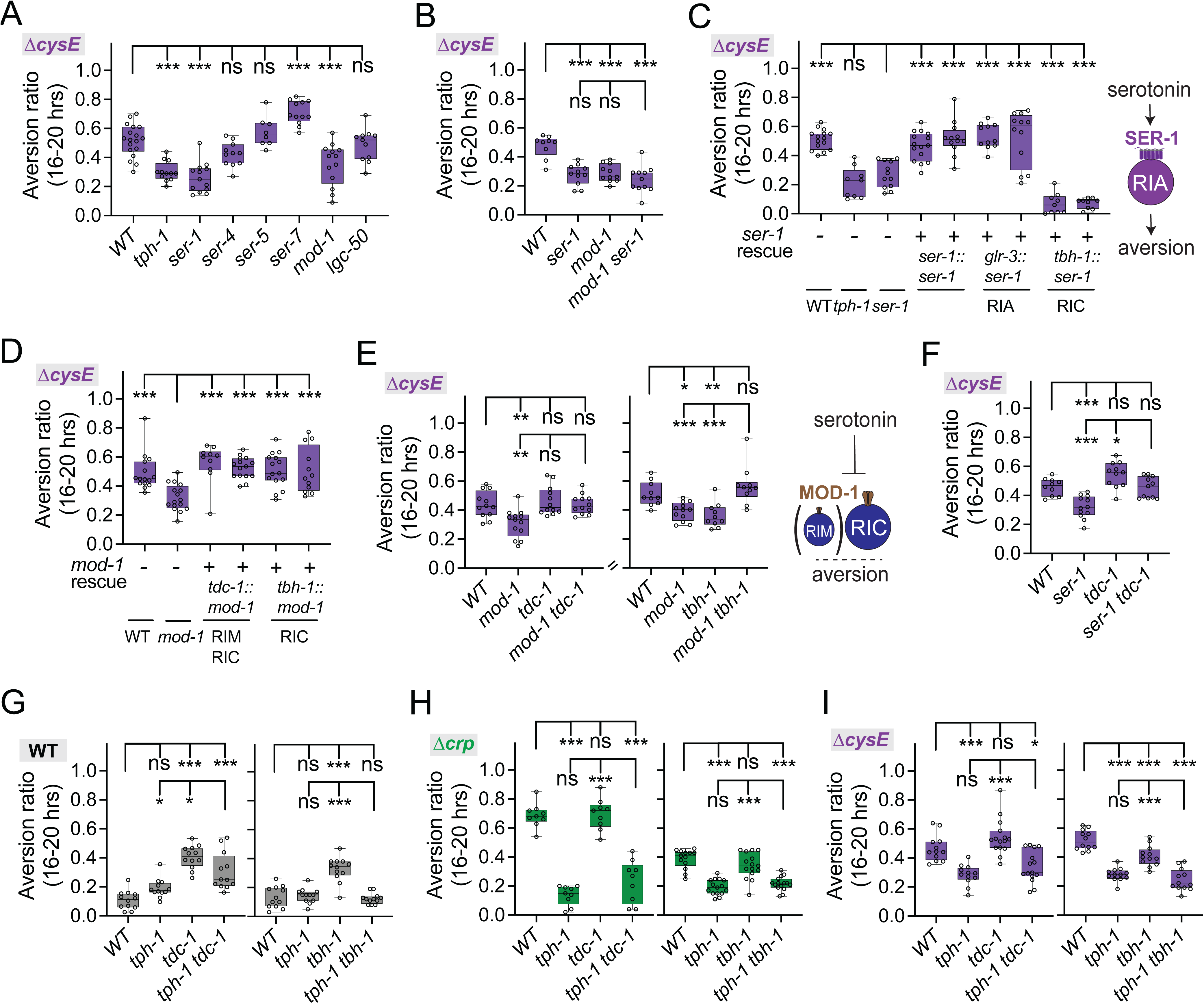
Aversion from the mediocre Δ*cysE* diet requires two serotonin receptors, SER-1 and MOD-1. (A) Behavioral responses of wild-type, *tph-1*, and serotonin receptor mutant animals on the mediocre Δ*cysE* diet. Aversion is diminished in *ser-1* and *mod-1* mutants. (B) Genetic interaction of *mod-1* with *ser-1* on the mediocre Δ*cysE* diet. (C) Aversion behavior of *ser-1* mutants expressing *ser-1* cDNA from its endogenous promoter (*ser-1)*, or promoters for RIA (*glr-3)*, or RIC neuron (*tbh-1*). Two transgenic lines were tested for each plasmid. (D) Aversion behavior of *mod-1* rescue in RIM and/or RIC neuron. Two transgenic lines were tested for each plasmid. (E) Genetic interaction of *mod-1* with *tdc-1* or *tbh-1* on the Δ*cysE* diet and schematic model of MOD-1 action. Stopped and dotted lines represent inhibition of RIC and octopamine. (F) Genetic interaction of *ser-1* with *tdc-1* on the mediocre Δ*cysE* diet. (G-I) Genetic interactions of *tph-1, tdc-1,* and *tbh-1* across bacterial diets. (G) wild-type diet; (H) Δ*crp* diet; (I) Δ*cysE* diet. For (A) to (F), each data point indicates individual assay. Results are shown with median ± quartiles in boxes and Min to Max whiskers. ns, not significant, *P<0.05, **P<0.01, ***P<0.001 by One-Way ANOVA, corrected by Dunnett’s multiple comparisons (panel A, C, D), or Tukey’s multiple comparisons (panel B, E-I).

Expression of the inhibitory MOD-1 serotonin receptor in RIC neurons or in RIM and RIC neurons rescued aversion of *mod-1* mutants on the mediocre Δ*cysE* diet (**Figure 7D**), in agreement with prior results showing that serotonin from NSM can inhibit RIM via MOD-1.^67^ Double mutants between the RIM/RIC neurotransmitter enzymes *tdc-1* or *tbh-1* and *mod-1* restored aversion on the mediocre Δ*cysE* diet (**Figure 7E**). The combination of *mod-1* rescue in RIC neurons and aversion in *mod-1 tbh-1* double mutants indicates that octopamine from RIC can suppress aversion from the Δ*cysE* diet, but is inhibited by *mod-1*.

Notably, different combinations of genes, neurons, and bacterial diets did not always yield simple interpretations. For example, *tdc-1 ser-1* double mutants had strong aversion to the Δ*cysE* diet, indicating that *tdc-1* can suppress *ser-1,* even though *ser-1* aversion was not rescued in the RIM and RIC *tdc-1-*expressing neurons (**Figure 7F**). The dopamine-deficient *cat-2* mutant had reduced aversion to the mediocre Δ*cysE* diet, but CEP neurons and the dopamine receptor *dop-6* were not specifically required for this behavior, further distinguishing the requirements for aversion from Δ*crp* and Δ*cysE* mediocre diets **(Figure S4A and S4B)**.

Finally, genetic interactions between *tph-1, tdc-1,* and *tbh-1* highlighted the context-dependent contributions of serotonin, tyramine, and octopamine on different bacterial diets (**Figure 7G-I**). Double mutants between *tph-1* and *tbh-1* resembled *tph-1* mutants on all diets, pointing to a central role of serotonin across conditions, and a secondary role for octopamine. However, on a wild-type diet tyramine may have at least some serotonin-independent functions, as *tdc-1 tph-1* double mutants had intermediate phenotypes compared to either single mutant (**Figure 7G**).

## DISCUSSION

### Aversion is a quantitative, serotonin-dependent foraging behavior

Ingestion of toxic foods that induce sickness, nausea, or gastrointestinal dysfunction is followed by aversion behaviors that prolongs an animal’s survival.^1,68^ After a few hours of exposure, *C. elegans* abandons lethal bacterial foods such as certain RNAi-expressing bacteria,^27^ food spiked with exogenous toxins,^27,57^ or pathogens.^36,38,69^ Here we show that a less extreme, but related aversion behavior is induced by a small number of mutant *E. coli* diets. This behavior is observed in contexts in which animals grow and reproduce normally, but express mitochondrial or redox stress markers. We suggest that aversion represents a quantitative behavioral response to mediocre food quality that reflects metabolic stress.

Our results identify a set of interacting, partly cooperative and partly antagonistic neurotransmitters and receptors that regulate aversion behavior. The neurotransmitter serotonin is required for *C. elegans* to distinguish between high-quality and mediocre *E. coli* diets, with bidirectional roles in guiding behavior (**Figure 8A** and **8B**). Serotonin has generally been associated with positive aspects of food quality in *C. elegans,* and stimulates behaviors such as feeding, egg-laying, and dwelling that are observed in beneficial conditions. However, serotonin has a prominent role in behavioral and physiological responses to toxic foods in mammals,^70^ and in *C. elegans,* serotonin is required for the avoidance of toxic or pathogenic bacteria,^27,57^ and for the learned avoidance of bacterial pathogens.^42^ Using cell-specific knockouts and acute silencing experiments, we found that high-quality and mediocre diets were interpreted by different classes of serotonergic neurons. The serotonergic ADF neurons supported retention on a high-quality bacterial diet, while serotonergic NSM neurons and to a lesser extent HSN neurons drove aversion from mediocre diets (**Figure 8B**). ADF is a chemosensory neuron that is activated by bacterial metabolites such as polyamines^43^, consistent with a role in assessing food quality. In previous studies, NSM has been most strongly implicated in sensing high-quality food,^41^ but NSM can also shift its properties to drive a food avoidance response after animals are exposed to food spiked with the mitochondrial toxin antimycin A.^71^ Our results suggest that mediocre diets may induce a related NSM-dependent aversive response.

**Figure 8.**
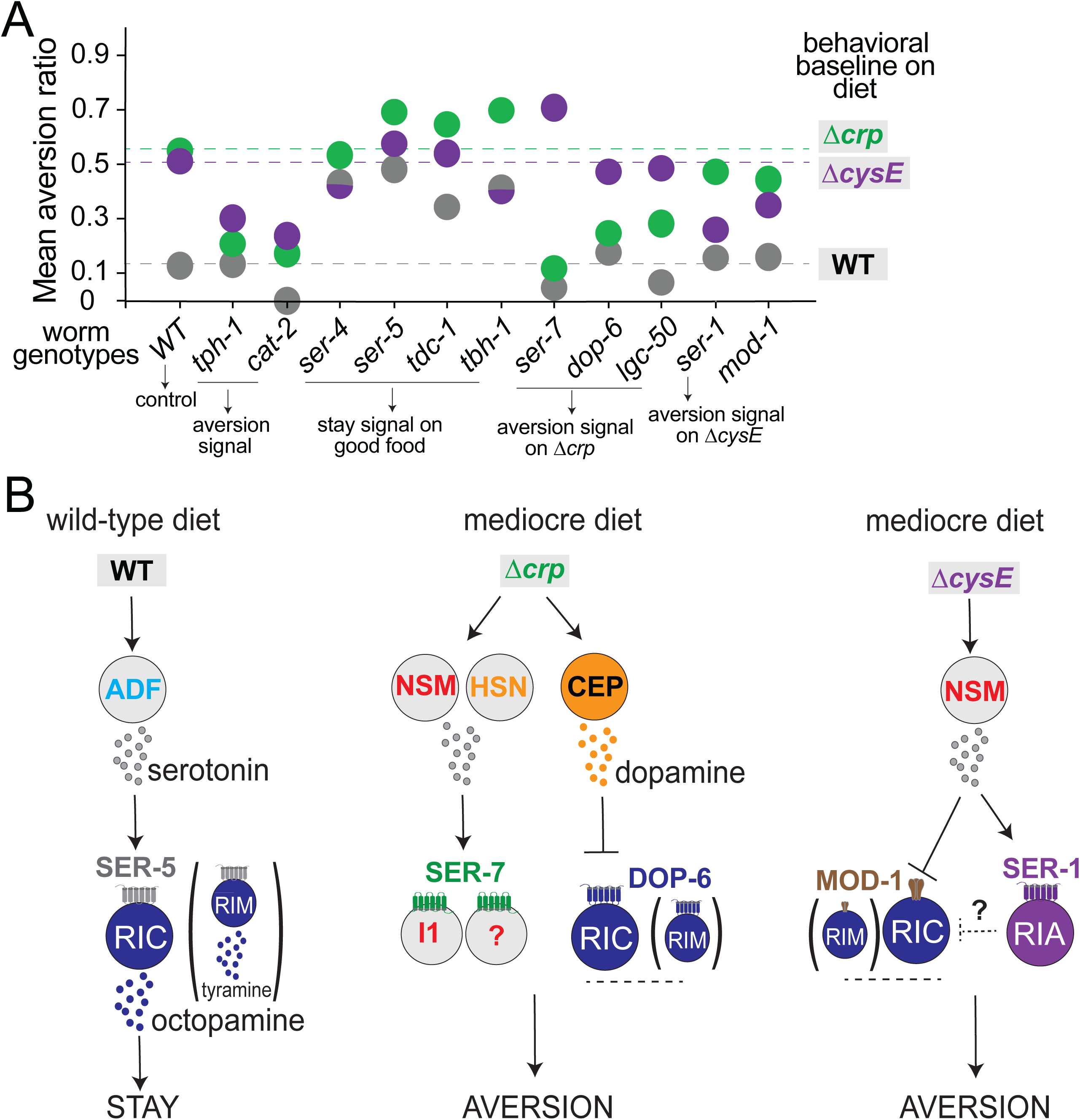
Summary and model of how *C. elegans* distinguishes food quality. (A) Neurotransmitters and receptors that regulate aversion on the wild-type diet and mediocre Δ*crp* or Δ*cysE* diets. Mean aversion ratios across all assays were used to generate the bubble map; dotted lines represent the mean aversion ratio of wild-type animals on each diet; statistical comparisons appear in the main figures. (B) Preliminary neural circuits for aversion behavior; additional neurons remain to be defined. Arrows represent activation of neurons, transmitters, or behaviors; stopped and dotted lines represent inhibition.

By screening for aversion behavior on three different bacterial diets, we identified roles for all six known serotonin receptors in the aversion assay. Two of the serotonin receptors, *ser-4* and *ser-5,* were required to suppress aversion from a high-quality diet; the other four receptors, *ser-7, lgc-50, mod-1,* and *ser-1,* supported aversion from two mediocre diets (**Figure 8A**). Some serotonin receptors distinguished among mediocre foods, such that *ser-7* only affected aversion to Δ*crp,* whereas *ser-1* only affected aversion to Δ*cysE*. This double dissociation indicates that the nervous system can discriminate among aversive contexts.

All six serotonin receptors also participate in a serotonin-dependent slowing behavior when *C. elegans* enters a bacterial lawn.^45^ Slowing on lawn entry relies on NSM and HSN serotonergic neurons, like aversion from mediocre diets, but the contributions of receptors are different: *mod-1, ser-4,* and *lgc-50* have aligned and partly redundant functions that promote slowing behavior, while *ser-1, ser-5,* and *ser-7* have modulatory roles.^45^ In another assay, food-dependent regulation of nociception, the serotonin receptors *ser-1, ser-5,* and *mod-1* have congruent but non-redundant roles.^72^ These comparisons highlight the distinctiveness of serotonin signaling in different contexts.

### Dopamine, octopamine, and tyramine regulate aversion behavior through specific neural circuits

In addition to serotonin, the monoamine transmitters dopamine and octopamine also contribute to aversion behavior. Dopamine promoted aversion from both mediocre diets, while octopamine suppressed aversion from all diets.

Octopamine and tyramine are considered the invertebrate counterparts of vertebrate norepinephrine and epinephrine; all of these transmitters are synthesized from the amino acid tyramine and signal through G protein-coupled receptors. We found that the serotonin and dopamine receptors *ser-5, mod-1,* and *dop-6* could all modulate aversion when expressed in the octopaminergic RIC neurons, alone or together with the tyraminergic RIM neurons. Our results suggest that on the wild-type diet, serotonin from the ADF chemosensory neurons acts via the excitatory SER-5 receptor to enhance the activity of RIC; on the mediocre Δ*crp* diet, dopamine from the CEP mechanosensory neurons acts via the DOP-6 receptor to inhibit RIC; and on the mediocre Δ*cysE* diet, serotonin from the NSM neurons acts via the serotonin-gated chloride channel MOD-1 to inhibit RIC.

Octopamine and tyramine also suppress aversion to the bacterial pathogen *Pseudomonas aeruginosa* PA14.^38^ Exposure to the pathogen downregulates expression of the biosynthetic enzyme *tdc-1* in RIM and RIC neurons, resulting in reduced tyramine and octopamine signaling.^38^ Together, these results implicate RIM and RIC neurons, and their tyramine and octopamine transmitters, in behavioral responses to multiple aversive conditions.

Across all *E. coli* diets and double mutant contexts, the effects of serotonin (*tph-1)* were stronger than the effects of dopamine, tyramine, or octopamine. These results indicate that the RIC and RIM transmitters are not the only essential targets of serotonin signaling. One additional serotonin target is the RIA neuron, which integrates sensory responses to regulate forward movement, and is a site of *ser-1* rescue on the mediocre Δ*cysE* diet. Another serotonin target is the I1 neuron, a little-studied neuron in the pharyngeal nervous system that is a site of *ser-7* rescue on the mediocre Δ*crp* diet. Tracing the interactions between these neurons and transmitters will allow further elucidation of these neuromodulatory circuits and their responses to different bacterial diets.

### Enriched metabolic pathways in bacteria affect animal physiology and behavior

Our genome-wide screen identified 22 *E. coli* mutants that induced aversive responses in *C. elegans*, which represent only a small fraction of the 244 *E. coli* mutants that result in *C. elegans* developmental delay.^14^ Aversion is a stringent assay that requires animals to spend a substantial fraction of their time away from the only available food; more sensitive behavioral assays such as choices between two bacteria would likely uncover more distinctions among bacterial strains.

We focused our studies on two bacterial genes whose effects on *C. elegans* had not been examined previously, the catabolite regulator CRP and the cysteine biosynthesis gene *cysE*. *crp* is a global regulator of bacterial metabolism on low-carbohydrate media. Under the growth conditions used here, Δ*crp* bacteria were large in size and induced expression of a stress reporter induced by toxic or inedible bacteria (*daf-*7) and a reporter for the mitochondrial unfolded protein response *(hsp-6)*. These two reporters were also induced by the Δ*fepB* ferric enterobactin transport mutant that elicited aversion. Interestingly, our screen yielded the same four *fes/fep* genes as screens for *C. elegans* developmental delay,^14^ poor development on a nutrient-limited food source,^73^ and synthetic lethality with the *C. elegans* mitochondrial mutant *spg-7*.^16^ The ferric-enterobactin complex that accumulates in these strains appears to cause mitochondrial toxicity.^16^ As mitochondrial toxins can drive food aversion in *C. elegans*;^27^ we suggest that mitochondrial dysfunction contributes to aversion to Δ*fes/fep* and Δ*crp* diets. Aversion to the mediocre Δ*crp* diet may also integrate a mechanosensory response to abnormal bacterial size or shape that makes the bacteria harder to ingest, which can lead to behavioral aversion.^37^

The identification of *cysE* as a mediocre food that elicits the oxidative stress reporter *gst-4* (i.e., glutathione S-transferase) is consistent with the role of cysteine in synthesis of the antioxidant glutathione. *C. elegans* synthesizes glutathione from cysteine and other precursors, and this pathway requires more cysteine than protein synthesis. Either an excess or a deficiency of dietary thiols has effects on redox states that are disadvantageous for *C. elegans*,^74^ explaining why cysteine, an amino acid that *C. elegans* can synthesize on its own, nonetheless can be limited by bacterial production and disruption of glutathione homeostasis.

The *C. elegans* stress responses induced by mediocre food, pathogens, and toxins overlap, as do the genes that regulate aversion. We suggest that the evaluation of food quality across different contexts converges on a shared, distributed circuit for survival-relevant aversion behaviors.

### Limitations of the study

We identified neurons in which specific neurotransmitter receptors can rescue behavior, but these may not represent all neurons that are necessary or sufficient for receptor function. In previous studies of the neuropeptide receptor gene *dmsr-7* in a pathogen aversion assay, several non-overlapping groups of cells that were each sufficient to rescue the *dmsr-7* mutant defect.^38^ The same principle could apply more broadly in neuromodulatory pathways.

While tyramine and octopamine are synthesized by RIM and RIC neurons, they are also synthesized by neuroendocrine cells in the somatic gonad, the ventral uterine cells (tyramine) and gonadal sheath cells (octopamine).^33^ The *tbh-1* promoter used for RIC expression is also expressed in gonadal sheath cells, and the *tdc-1* promoter used for RIM and RIC expression is also expressed in ventral uterine cells. The intersection of endogenous expression and promoters used for rescue experiments point to RIC as the key site of *ser-5, dop-6,* and *mod-1* action, but it is possible that octopamine from the gonadal sheath cells also has a role.^75,76^

Finally, these studies do not include direct functional measurements of neuronal activity under different conditions. A variety of functional changes, including changes in excitability, neurotransmitter release, or gene expression would be consistent with the activating or inhibitory interactions inferred from these genetic interactions.

## STAR★METHODS

### KEY RESOURCES TABLE

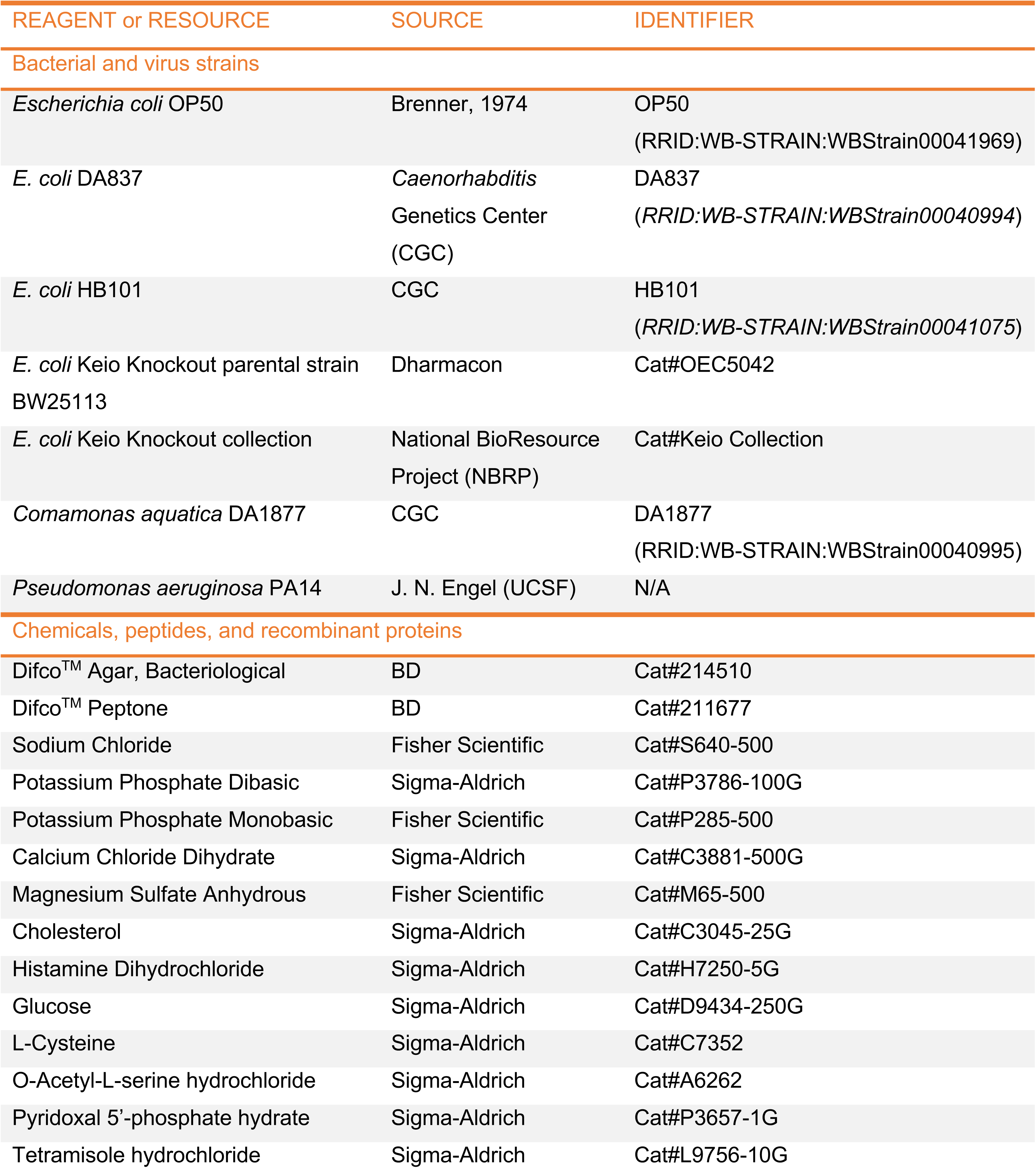
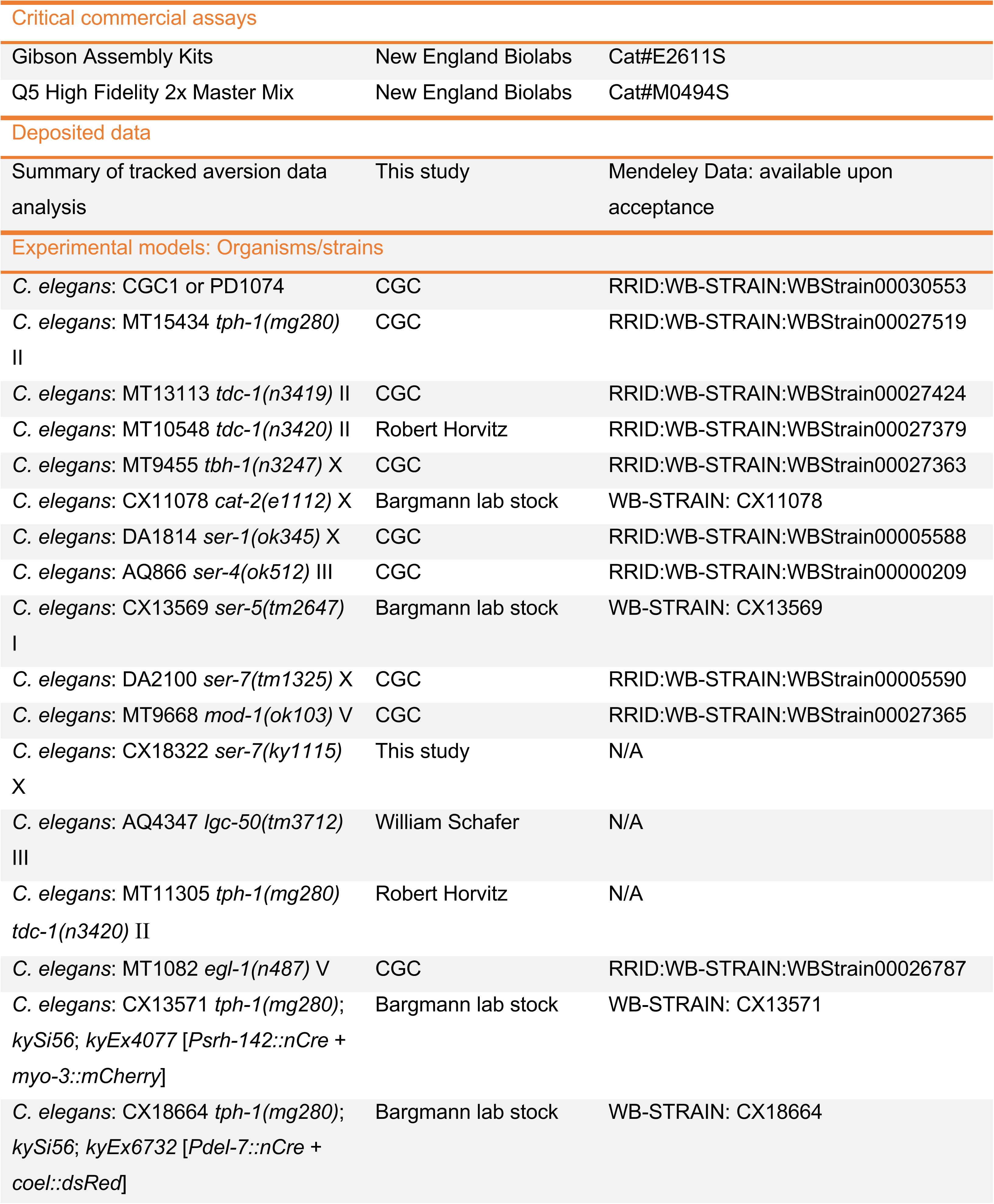
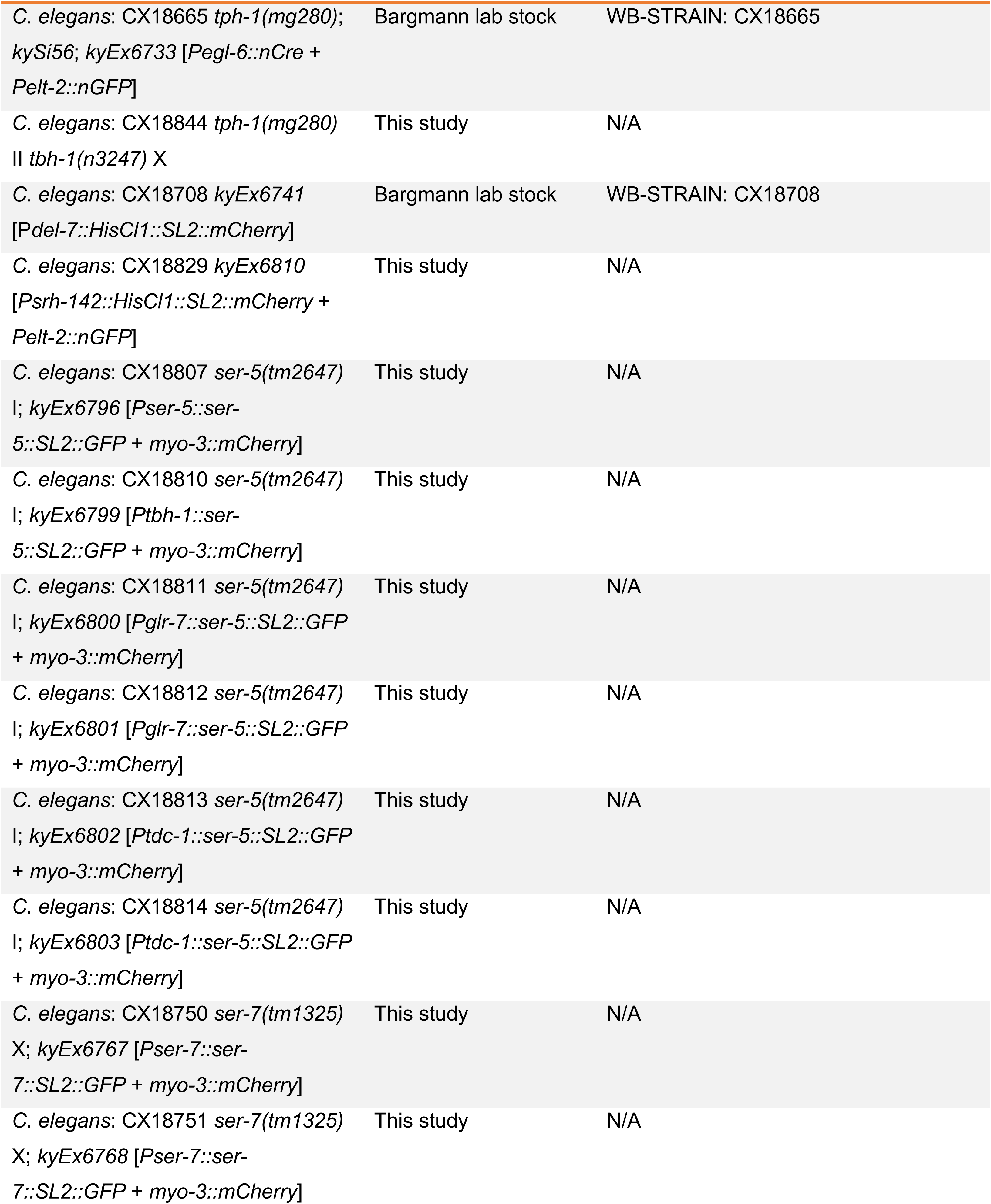
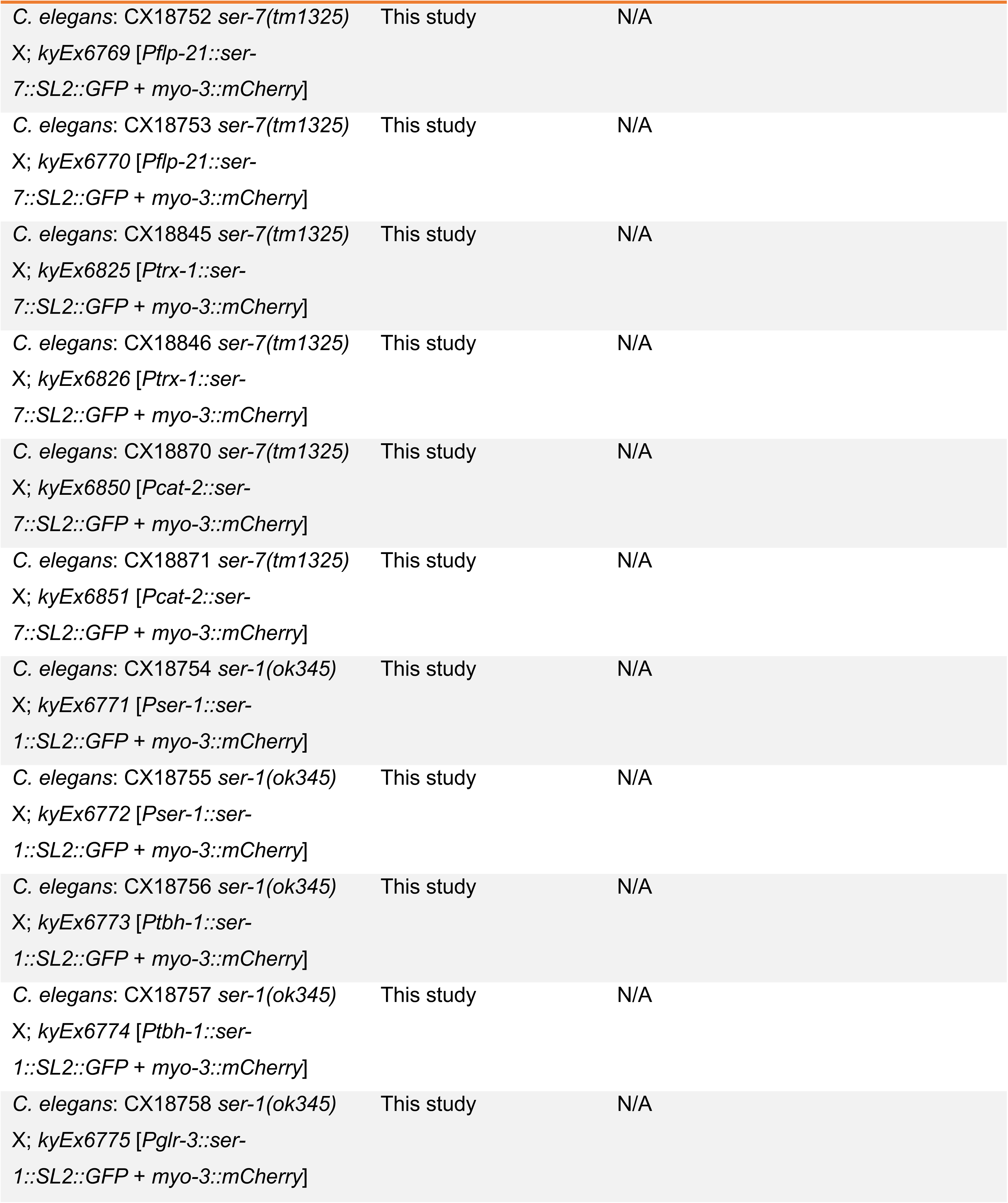
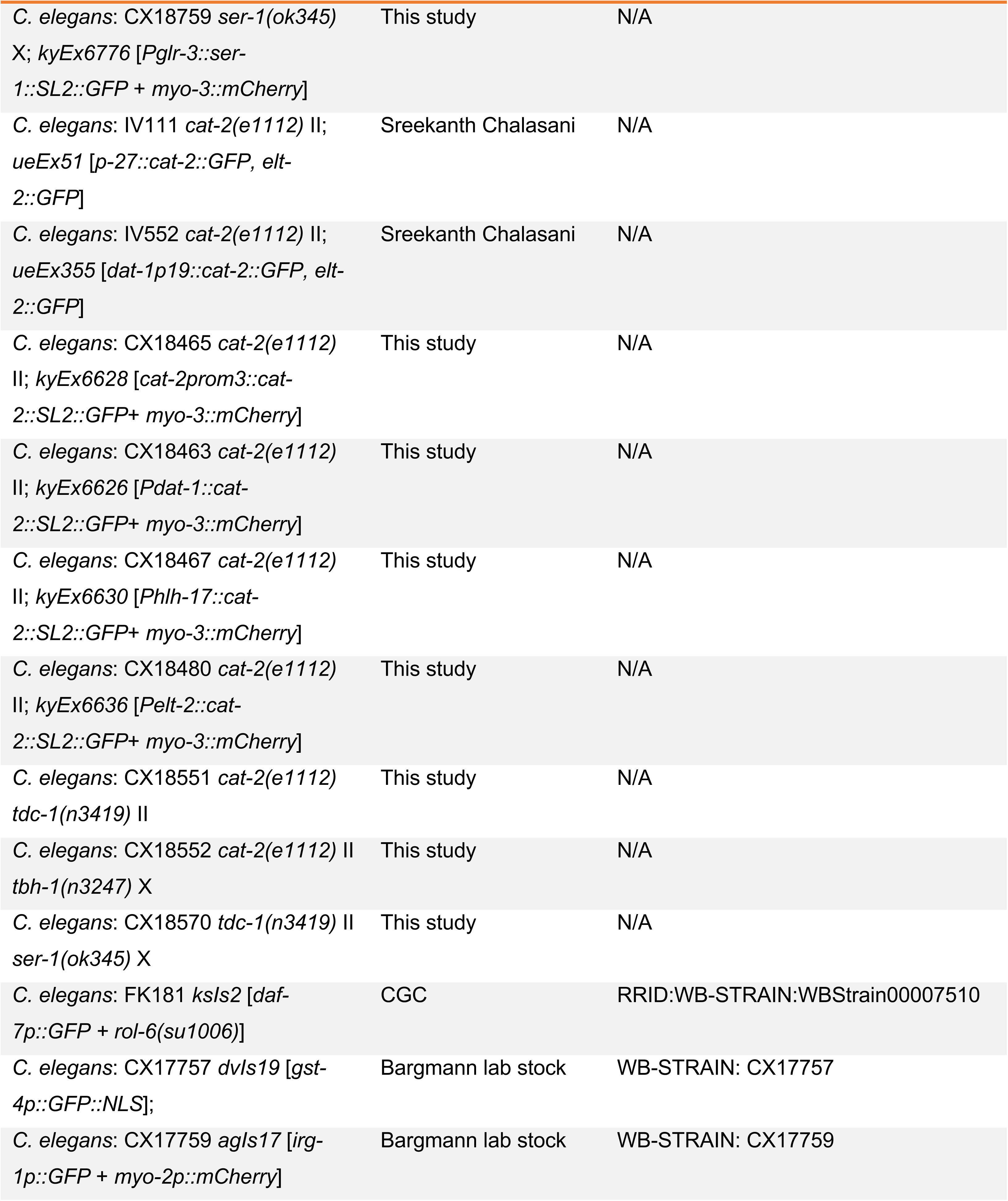
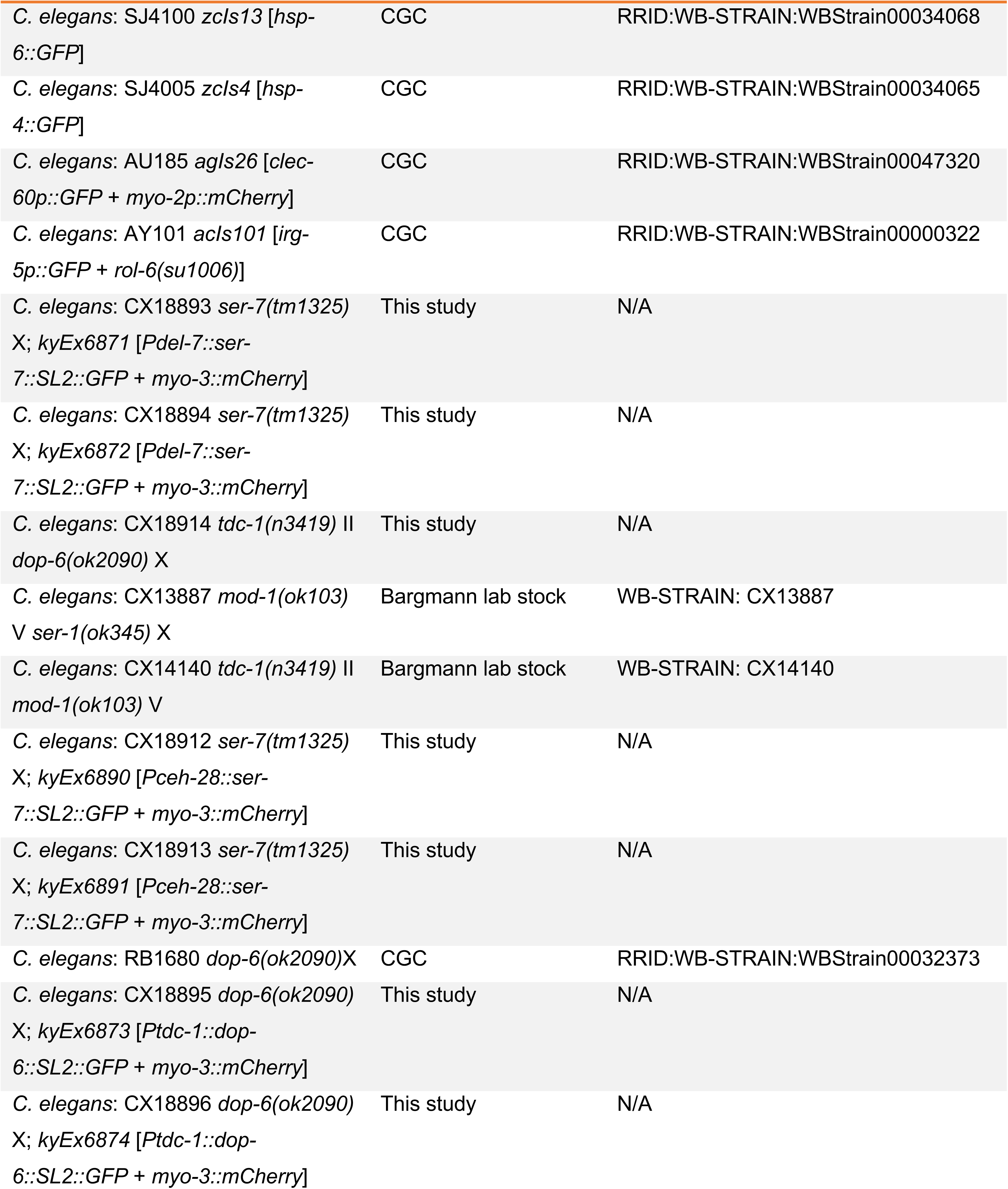
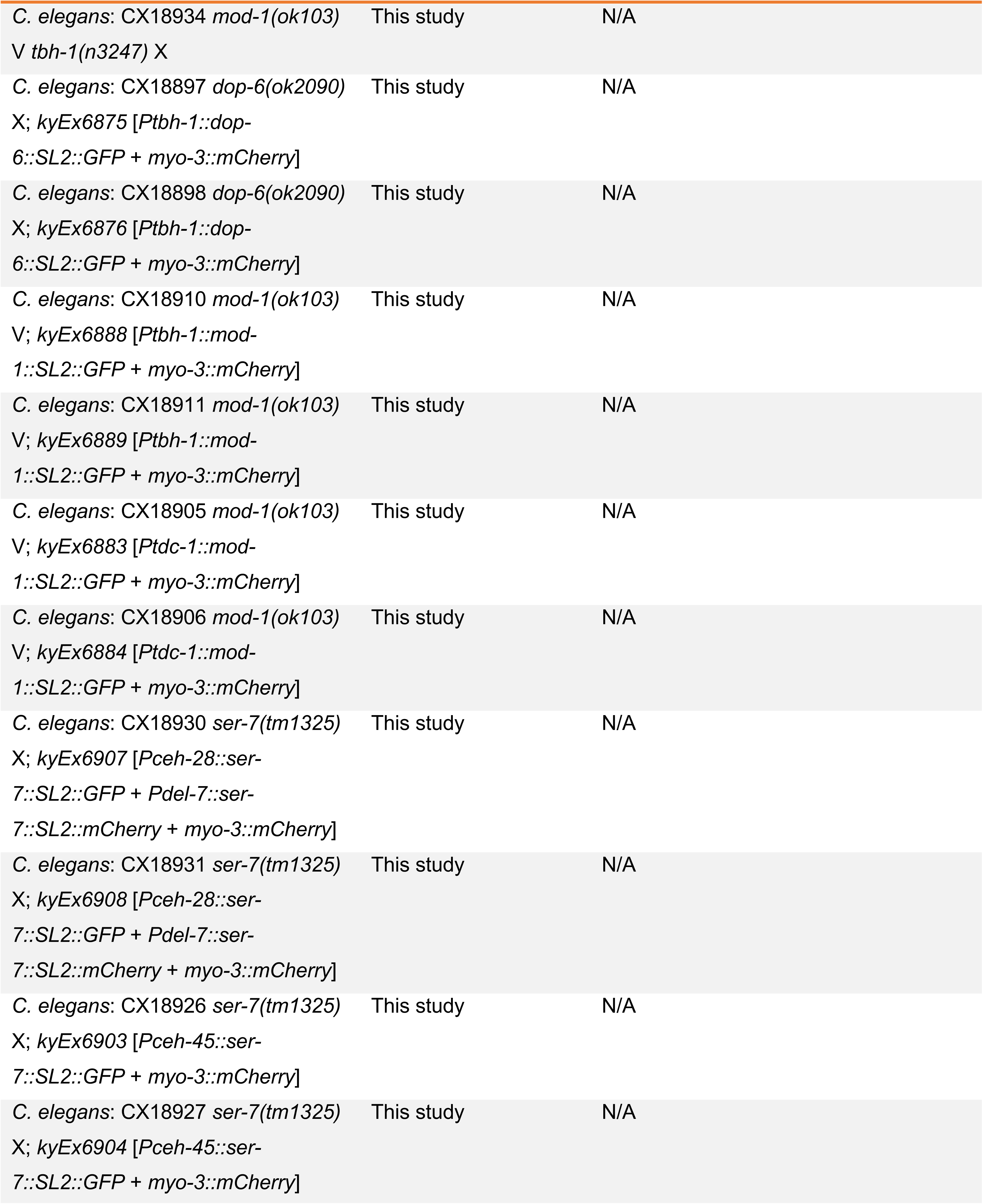
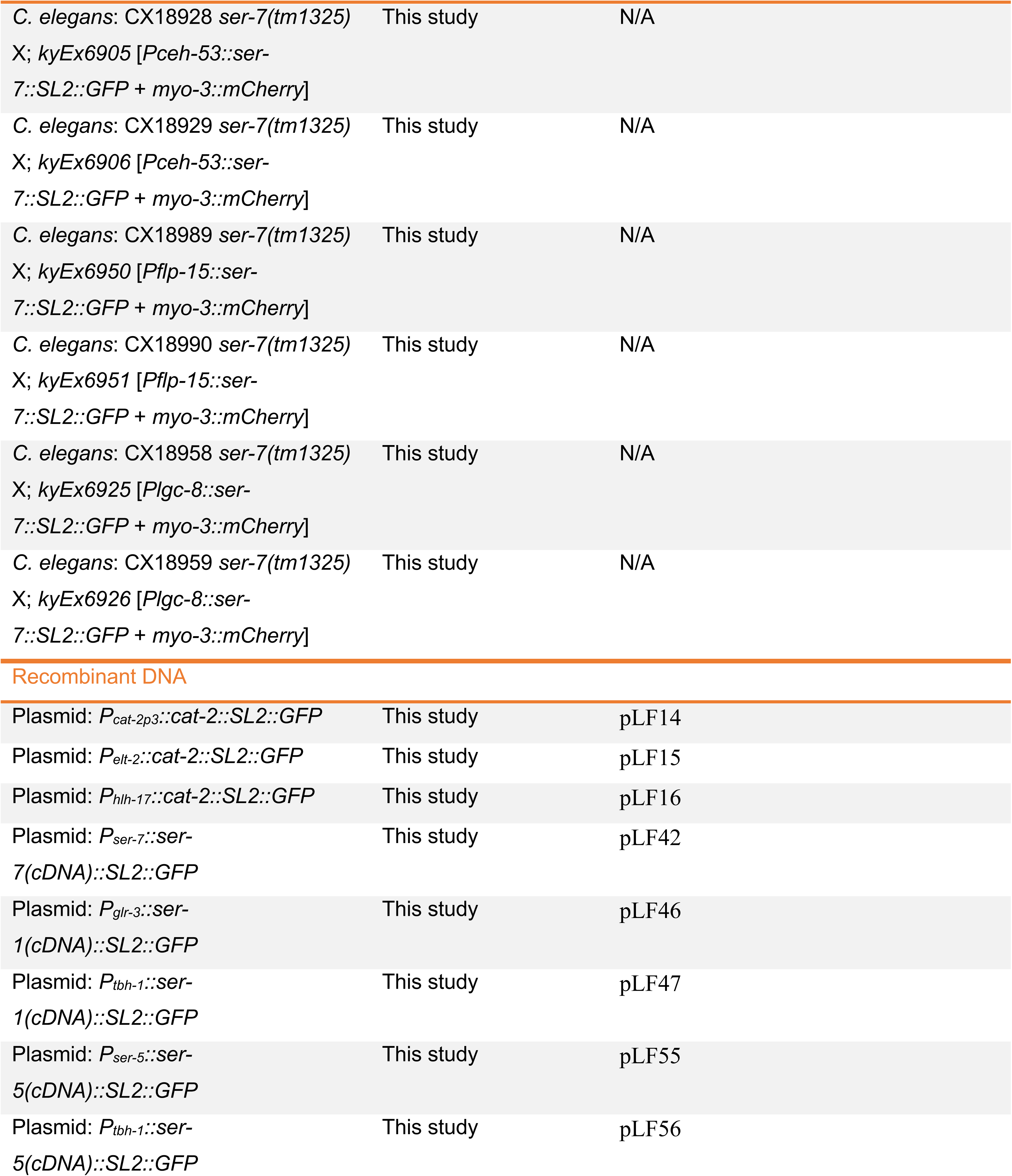
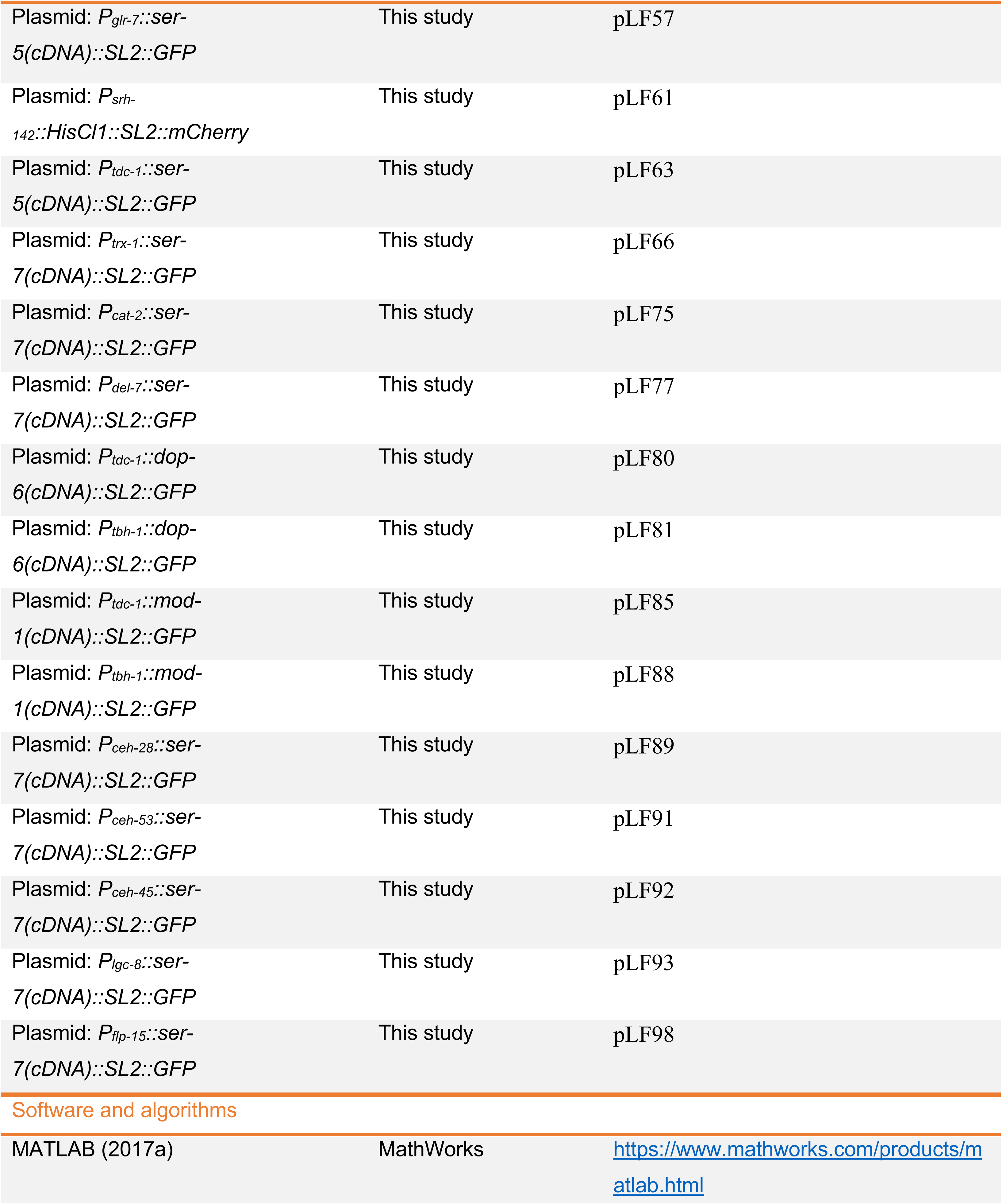
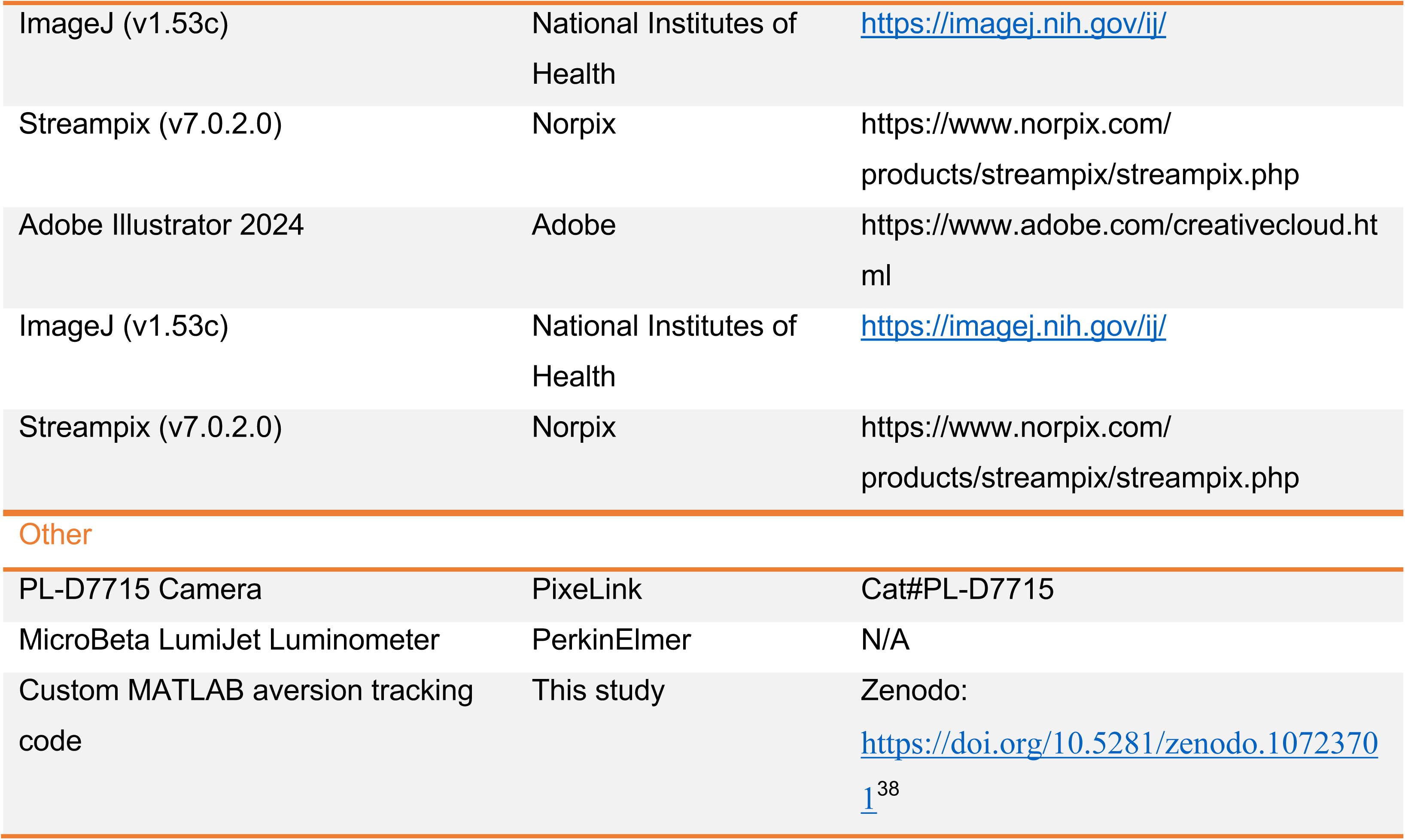

### RESOURCE AVAILABILITY

#### Lead contact

All information and requests for reagents or strains should be directed to the lead contact Cori Bargmann.

#### Materials availability

All reagents and strains used in this study are available upon request to the lead contact.

## EXPERIMENTAL MODEL AND SUBJECT DETAILS

### C. elegans strains

All *C. elegans* strains were maintained on nematode growth medium (NGM) agar plates seeded with bacterial diet *E. coli* BW25113 at 20 °C. CGC1 (formerly known as PD1074) was provided by the Caenorhabditis Genetics Center (CGC) and used as the wild-type strain. The other strains obtained from the CGC are: MT15434 *tph-1(mg280)*; MT13113 *tdc-1(n3419)*; MT9455 *tbh-1(n3247)*; MT1082 *egl-1(n487*); DA1814 *ser-1(ok345)*; AQ866 *ser-4(ok512)*; DA2100 *ser-7(tm1325)*; MT9668 *mod-1(ok103)*; RB1680 *dop-6(ok2090)*; FK181 *ksIs2* [*daf-7p::GFP* + *rol-6(su1006)*]; SJ4100 *zcIs13* [*hsp-6::GFP*]; SJ4005 *zcIs4* [*hsp-4::GFP*]; AU185 *agIs26* [*clec-60p::GFP* + *myo-2p::mCherry*]; AY101 *acIs101* [*irg-5p::GFP* + *rol-6(su1006)*]

The following strains were obtained from the Bargmann strain collection: CX11078 *cat-2(e1112)*; CX13571 *tph-1(mg280)*; *kySi56* [single-copy mosSCI insertion of floxed tph-1(genomic) sequence];^25^ *kyEx4077* [*Psrh-142::nCre* + *myo-3::mCherry*]; CX18664 *tph-1(mg280)*; *kySi56*; *kyEx6732* [*Pdel-7::nCre* + *coel::dsRed*]; CX18665 *tph-1(mg280)*; *kySi56*; *kyEx6733* [*Pegl-6::nCre* + *Pelt-2::nGFP*]; CX13569 *ser-5(tm2647)*; CX18322 *ser-7(ky1115)*; CX18708 *kyEx6741* [P*del-7::HisCl1::SL2::mCherry*]; CX17757 *dvIs19* [*gst-4p::GFP::NLS*]; CX17759 *agIs17* [*irg-1p::GFP* + *myo-2p::mCherry*]; CX13887 *mod-1(ok103)* V *ser-1(ok345)* X; CX14140 *tdc-1(n3419)* II *mod-1(ok103)* V; CX14993 *kyEx4962* [*tdc-1::TeTx::mCherry*]; CX17912 *kyEx6345* [*tbh-1::TeTx::mCherry + coel::GFP*] The following strains were obtained from Dr. Robert Horvitz^33^: MT10661 *tdc-1(n3420)*; MT11305 *tph-1(mg280) tdc-1(n3420)* The following strain was obtained from Dr. William Schafer^77^: AQ4347 *lgc-50(tm3712)* The following strains were obtained from Dr. Sreekanth Chalasani^78^: IV111 *cat-2(e1112)* II; *ueEx51* [*p-27::cat-2::GFP, elt-2::GFP*]; IV552 *cat-2(e1112)* II; *ueEx355* [*dat-1p19::cat-2::GFP, elt-2::GFP*]

The following strains were generated in this study: CX18844 *tph-1(mg280)* II *tbh-1(n3247)* X CX18551 *cat-2(e1112) tdc-1(n3419)* II CX18552 *cat-2(e1112)* II *tbh-1(n3247)* X CX18570 *tdc-1(n3419)* II *ser-1(ok345)* X CX18914 *tdc-1(n3419)* II *dop-6(ok2090)* X CX18934 *mod-1(ok103)* V *tbh-1(n3247)* X CX18829 *kyEx6810* [*Psrh-142::HisCl1::SL2::mCherry* + *Pelt-2::nGFP*]

– HisCl1 expressing in ADF, ADF silencing in the presence of histamine. CX18807 *ser-5(tm2647)* I; *kyEx6796* [*Pser-5::ser-5::SL2::GFP* + *myo-3::mCherry*]
– *ser-5* full rescue, line A. CX18808 *ser-5(tm2647)* I; *kyEx6797* [*Pser-5::ser-5::SL2::GFP* + *myo-3::mCherry*]
– *ser-5* full rescue, line B. CX18809 *ser-5(tm2647)* I; *kyEx6798* [*Ptbh-1::ser-5::SL2::GFP* + *myo-3::mCherry*]
– *ser-5* rescue in RIC neuron and gonadal sheath cells, line A. CX18810 *ser-5(tm2647)* I; *kyEx6799* [*Ptbh-1::ser-5::SL2::GFP* + *myo-3::mCherry*]
– *ser-5* rescue in RIC neuron and gonadal sheath cells, line B. CX18811 *ser-5(tm2647)* I; *kyEx6800* [*Pglr-7::ser-5::SL2::GFP* + *myo-3::mCherry*]
– *ser-5* rescue in pharyngeal neurons (I3, I6, *etc*), line A. CX18812 *ser-5(tm2647)* I; *kyEx6801* [*Pglr-7::ser-5::SL2::GFP* + *myo-3::mCherry*]
– *ser-5* rescue in pharyngeal neurons (I3, I6, *etc*), line B. CX18813 *ser-5(tm2647)* I; *kyEx6802* [*Ptdc-1::ser-5::SL2::GFP* + *myo-3::mCherry*]
– *ser-5* rescue in RIM, RIC and germline, line A. CX18814 *ser-5(tm2647)* I; *kyEx6803* [*Ptdc-1::ser-5::SL2::GFP* + *myo-3::mCherry*]
– *ser-5* rescue in RIM, RIC and germline, line B. CX18750 *ser-7(tm1325)* X; *kyEx6767* [*Pser-7::ser-7::SL2::GFP* + *myo-3::mCherry*]
– *ser-7* full rescue, line A. CX18751 *ser-7(tm1325)* X; *kyEx6768* [*Pser-7::ser-7::SL2::GFP* + *myo-3::mCherry*]
– *ser-7* full rescue, line B. CX18752 *ser-7(tm1325)* X; *kyEx6769* [*Pflp-21::ser-7::SL2::GFP* + *myo-3::mCherry*]
– *ser-7* rescue in pharyngeal neurons (M4, MC, M2, *etc*), line A. CX18753 *ser-7(tm1325)* X; *kyEx6770* [*Pflp-21::ser-7::SL2::GFP* + *myo-3::mCherry*] – *ser-7* rescue in pharyngeal neurons (M4, MC, M2, *etc*), line B.

CX18845 *ser-7(tm1325)* X; *kyEx6825* [*Ptrx-1::ser-7::SL2::GFP* + *myo-3::mCherry*] – *ser-7* rescue in ASJ, line A.

CX18846 *ser-7(tm1325)* X; *kyEx6826* [*Ptrx-1::ser-7::SL2::GFP* + *myo-3::mCherry*] – *ser-7* rescue in ASJ, line B.

CX18893 *ser-7(tm1325)* X; *kyEx6871* [*Pdel-7::ser-7::SL2::GFP* + *myo-3::mCherry*] – *ser-7* rescue in NSM, line A.

CX18894 *ser-7(tm1325)* X; *kyEx6872* [*Pdel-7::ser-7::SL2::GFP* + *myo-3::mCherry*] – *ser-7* rescue in NSM, line B.

CX18912 *ser-7(tm1325)* X; *kyEx6890* [*Pceh-28::ser-7::SL2::GFP* + *myo-3::mCherry*] – *ser-7* rescue in M4, line A.

CX18913 *ser-7(tm1325)* X; *kyEx6891* [*Pceh-28::ser-7::SL2::GFP* + *myo-3::mCherry*] – *ser-7* rescue in M4, line B.

CX18930 *ser-7(tm1325)* X; *kyEx6907* [*Pceh-28::ser-7::SL2::GFP* + *Pdel-7::ser-7::SL2::mCherry* + *myo-3::mCherry*] – *ser-7* rescue in M4 and NSM, line A. CX18931 *ser-7(tm1325)* X; *kyEx6908* [*Pceh-28::ser-7::SL2::GFP* + *Pdel-7::ser-7::SL2::mCherry* + *myo-3::mCherry*] – *ser-7* rescue in M4 and NSM, line B. CX18926 *ser-7(tm1325)* X; *kyEx6903* [*Pceh-45::ser-7::SL2::GFP* + *myo-3::mCherry*]

– ser-7 rescue in MI, I1 and I3, line A. CX18927 ser-7(tm1325) X; kyEx6904 [Pceh-45::ser-7::SL2::GFP + myo-3::mCherry]
– ser-7 rescue in MI, I1 and I3, line B. CX18928 ser-7(tm1325) X; kyEx6905 [Pceh-53::ser-7::SL2::GFP + myo-3::mCherry]
– ser-7 rescue in M4 and I5, line A. CX18929 ser-7(tm1325) X; kyEx6906 [Pceh-53::ser-7::SL2::GFP + myo-3::mCherry]
– ser-7 rescue in M4 and I5, line B. CX18958 ser-7(tm1325) X; kyEx6925 [Plgc-8::ser-7::SL2::GFP + myo-3::mCherry]
– ser-7 rescue in I1, line B. CX18959 ser-7(tm1325) X; kyEx6926 [Plgc-8::ser-7::SL2::GFP + myo-3::mCherry]
– ser-7 rescue in I1, line C. CX18989 ser-7(tm1325) X; kyEx6950 [Pflp-15::ser-7::SL2::GFP + myo-3::mCherry]
– ser-7 rescue in I2, line A. CX18990 ser-7(tm1325) X; kyEx6951 [Pflp-15::ser-7::SL2::GFP + myo-3::mCherry]
– ser-7 rescue in I2, line B. CX18870 ser-7(tm1325) X; kyEx6850 [Pcat-2::ser-7::SL2::GFP + myo-3::mCherry]
– ser-7 rescue in dopaminergic neurons CEP, ADE, PDE, line A. CX18871 ser-7(tm1325) X; kyEx6851 [Pcat-2::ser-7::SL2::GFP + myo-3::mCherry]
– ser-7 rescue in dopaminergic neurons CEP, ADE, PDE, line B. CX18754 ser-1(ok345) X; kyEx6771 [Pser-1::ser-1::SL2::GFP + myo-3::mCherry]
– ser-1 full rescue, line A. CX18755 ser-1(ok345) X; kyEx6772 [Pser-1::ser-1::SL2::GFP + myo-3::mCherry]
– ser-1 full rescue, line B. CX18756 ser-1(ok345) X; kyEx6773 [Ptbh-1::ser-1::SL2::GFP + myo-3::mCherry]
– ser-1 rescue in RIC neuron and gonadal sheath cells, line A. CX18757 ser-1(ok345) X; kyEx6774 [Ptbh-1::ser-1::SL2::GFP + myo-3::mCherry]
– ser-1 rescue in RIC neuron and gonadal sheath cells, line B. CX18758 ser-1(ok345) X; kyEx6775 [Pglr-3::ser-1::SL2::GFP + myo-3::mCherry]
– ser-1 rescue in RIA neuron, line A. CX18759 ser-1(ok345) X; kyEx6776 [Pglr-3::ser-1::SL2::GFP + myo-3::mCherry]
– ser-1 rescue in RIA neuron, line B. CX18465 cat-2(e1112) II; kyEx6628 [cat-2prom3::cat-2::SL2::GFP+ myo-3::mCherry]
– cat-2 rescue in dopaminergic neurons CEP, ADE, PDE. CX18463 cat-2(e1112) II; kyEx6626 [Pdat-1::cat-2::SL2::GFP+ myo-3::mCherry]
– cat-2 rescue in dopaminergic neurons CEP, ADE, PDE. CX18467 cat-2(e1112) II; kyEx6630 [Phlh-17::cat-2::SL2::GFP+ myo-3::mCherry]
– cat-2 rescue in dopaminergic neurons CEP glia. CX18480 cat-2(e1112) II; kyEx6636 [Pelt-2::cat-2::SL2::GFP+ myo-3::mCherry]
– cat-2 rescue in intestine. CX18895 dop-6(ok2090) X; kyEx6873 [Ptdc-1::dop-6::SL2::GFP + myo-3::mCherry]
– dop-6 rescue in RIM, RIC neuron and germline, line A. CX18896 dop-6(ok2090) X; kyEx6874 [Ptdc-1::dop-6::SL2::GFP + myo-3::mCherry]
– dop-6 rescue in RIM, RIC neuron and germline, line B.

CX18897 *dop-6(ok2090)* X; *kyEx6875* [*Ptbh-1::dop-6::SL2::GFP* + *myo-3::mCherry*] – *dop-6* rescue in RIC neuron and gonadal sheath cells, line A.

CX18898 *dop-6(ok2090)* X; *kyEx6876* [*Ptbh-1::dop-6::SL2::GFP* + *myo-3::mCherry*] – *dop-6* rescue in RIC neuron and gonadal sheath cells, line B.

CX18910 *mod-1(ok103)* V; *kyEx6888* [*Ptbh-1::mod-1::SL2::GFP* + *myo-3::mCherry*] – *mod-1* rescue in RIC neuron and gonadal sheath cells, line A.

CX18911 *mod-1(ok103)* V; *kyEx6889* [*Ptbh-1::mod-1::SL2::GFP* + *myo-3::mCherry*] – *mod-1* rescue in RIC neuron and gonadal sheath cells, line B.

CX18905 *mod-1(ok103)* V; *kyEx6883* [*Ptdc-1::mod-1::SL2::GFP* + *myo-3::mCherry*] – *mod-1* rescue in RIM, RIC neuron and germline, line A.

CX18906 *mod-1(ok103)* V; *kyEx6884* [*Ptdc-1::mod-1::SL2::GFP* + *myo-3::mCherry*] – *mod-1* rescue in RIM, RIC neuron and germline, line B.

### Bacterial strains

*E. coli* strains OP50, DA837, HB101, BW25113, and *Comamonas aquatica* strain DA1877 were cultured in Lysogeny Broth (LB) liquid medium at 37 °C. *Pseudomonas aeruginosa* strain PA14 were cultured in Lysogeny Broth (LB) liquid medium at 25 °C. *E. coli* Keio deletion mutants^48^ were purchased from the National Bioresource Project of Japan (NBRP) and were grown at 37°C in LB medium with 25 μg/mL kanamycin.

## METHOD DETAILS

### Generation of transgenes

To construct plasmids for expression of *ser-5* in *C. elegans* neurons, promoter sequences of *ser-5* (2001bp), *glr-7* (2000bp), *tdc-1* (4550bp) or *tbh-1* (4537bp) were inserted into the *pSM::SL2::GFP::short unc-54 3’ UTR* vector along with a *ser-5* cDNA. Injection mixtures containing *Pser-5::ser-5::SL2::GFP* (30 ng/uL), *Pglr-7::ser-5::SL2::GFP* (30 ng/uL), *Ptdc-1::ser-5::SL2::GFP* (30 ng/uL), or *Ptbh-1::ser-5::SL2::GFP* (30 ng/uL) together with the co-injection marker *Pmyo-3::mCherry* (3 ng/uL) were injected into CX13569 *ser-5 (tm2647)* I.

To construct plasmids for expression of *ser-7* in *C. elegans* neurons, promoter sequences of *ser-7* (2118bp), *flp-21* (2003bp), *del-7* (839bp), *ceh-28* (1981bp), *ceh-53* (1476bp), *ceh-45* (2022bp), *trx-1* (1019bp), *lgc-8* (1999bp), *flp-15* (2402 bp) or *cat-2* (2137bp) were inserted into the *pSM::SL2::GFP::short unc-54 3’ UTR* vector along with a *ser-7* cDNA. Injection mixture containing *Pser-7::ser-7::SL2::GFP* (30 ng/uL), *Pflp-21::ser-7::SL2::GFP* (30 ng/uL), *Pdel-7::ser-7::SL2::GFP* (30 ng/uL), *Pceh-28::ser-7::SL2::GFP* (30 ng/uL), *Pceh-53::ser-7::SL2::GFP* (30 ng/uL), *Pceh-45::ser-7::SL2::GFP* (30 ng/uL), *Ptrx-1::ser-7::SL2::GFP* (30 ng/uL), *Plgc-8::ser-7::SL2::GFP* (30 ng/uL), *Pflp-15::ser-7::SL2::GFP* (30 ng/uL) or *Pcat-2::ser-7::SL2::GFP* (30 ng/uL) together with the co-injection marker *Pmyo-3::mCherry* (3 ng/uL) were injected into DA2100 *ser-7 (tm1325)* X.

To construct plasmids for expression of *ser-1* in *C. elegans* neurons, promoter sequences of *ser-1* (2103bp), *tbh-1* (4537bp), or *glr-3* (2838bp) were inserted into the *pSM::SL2::GFP::short unc-54 3’ UTR* vector along with a *ser-1* cDNA. Injection mixture containing *Pser-1::ser-1::SL2::GFP* (30 ng/uL), *Ptbh-1::ser-1::SL2::GFP* (30 ng/uL), or *Pglr-3::ser-1::SL2::GFP* (30 ng/uL) together with the co-injection marker *Pmyo-3::mCherry* (3 ng/uL) were injected into DA1814 *ser-1 (ok345)* X.

To construct plasmids for expression of histamine-gated chloride channel in order to chemically silencing *C. elegans* neurons, promoter sequence of *srh-142* (3976bp) was inserted into the *pSM::SL2::mCherry::short unc-54 3’ UTR* vector along with the HisCl1 coding sequence. Injection mixture containing *Psrh-142::HisCl1::SL2::mCherry* (20 ng/uL) together with the co-injection marker *Pelt-2::NLS::GFP* (2.5 ng/uL) were injected into PD1074.

To construct plasmids for expression of *cat-2* in *C. elegans* neurons, promoter sequences of *cat-2prom3* (1143bp), *dat-1* (786bp), *hlh-17* (3008bp), or *elt-2* (5049bp) were inserted into the *pSM::SL2::GFP:: unc-54 3’ UTR* vector along with the *cat-2* genomic coding region. Injection mixture containing *Pcat-2prom3::cat-2::SL2::GFP* (30 ng/uL), *Pdat-1::cat-2::SL2::GFP* (30 ng/uL), *Phlh-17::cat-2::SL2::GFP* (30 ng/uL), or *Pelt-2::cat-2::SL2::GFP* (30 ng/uL) together with the co-injection marker *Pmyo-3::mCherry* (5 ng/uL) were injected into CX11078 *cat-2 (e1112)* II.

To construct plasmids for expression of *dop-6* in *C. elegans* neurons, promoter sequences of *tdc-1* (4550bp) or *tbh-1* (4537bp) were inserted into the *pSM::SL2::GFP::short unc-54 3’ UTR* vector along with a *dop-6* cDNA. Injection mixture containing *Ptdc-1::dop-6::SL2::GFP* (30 ng/uL), or *Ptbh-1::dop-6::SL2::GFP* (30 ng/uL) together with the co-injection marker *Pmyo-3::mCherry* (3 ng/uL) were injected into RB1680 *dop-6 (ok2090)* X.

To construct plasmids for expression of *mod-1* in *C. elegans* neurons, promoter sequences of *tdc-1* (4550bp) or *tbh-1* (4537bp) were inserted into the *pSM::SL2::GFP::short unc-54 3’ UTR* vector along with a *mod-1* cDNA. Injection mixture containing *Ptdc-1::mod-1::SL2::GFP* (30 ng/uL), or *Ptbh-1::mod-1::SL2::GFP* (30 ng/uL) together with the co-injection marker *Pmyo-3::mCherry* (3 ng/uL) were injected into MT9668 *mod-1 (ok103)* V.

### Screen for *C. elegans* aversion to *E. coli* Keio knockout mutants

To prepare bacteria for the primary behavioral assay, individual *E. coli* deletion mutants were grown in LB medium with 25μg/mL kanamycin at 37°C for about 16 hours. 12.5μL overnight cultures were seeded onto 12-well plates containing 3.5 mL standard NGM agar in each well, dried, grown at 37°C for one day and incubated at room temperature for another day. Animals that were maintained on the standard wild-type bacteria BW25113 for at least two generations were synchronized for the behavioral assay. About 15-20 L4 animals were added onto each well of 12-well plates that contained BW25113_WT or individual *E. coli* mutants, and incubated at 21°C for 20 hours. Plater were then visually screened for the fraction of animals that exhibited aversive behavior, defined as (animals outside the bacterial lawn)/(total number of animals).

Each *E. coli* deletion mutant was screened for at least twice. A secondary screen was performed for identified positive hints from primary screen, using the same method. Positive hints identified from secondary screen were further verified in assays on 6-well plates that were video-tracked for 20 hours (recording rate 1 frame per minute, 1200 frames per video) with at least three independent replicates. Behavioral rigs were equipped with four 15 MP cameras (PL-D7715, Pixelink), one per six-well plate, and videos were cropped to individual wells and analyzed every six frames (200 data points per condition) using published MatLab codes (https://doi.org/10.5281/zenodo.10723701).^38^ Wild-type *C. elegans* on wild-type BW25113 bacteria were defined as having low aversive behavior (aversion ratio less than 0.2). Bacterial mutants that induced significantly higher aversion than BW25113_WT were chosen for subsequent studies.

### Genotyping identified *E. coli* mutants

Bacterial colonies from individual bacterial mutants were streaked on LB agar plates, grown at 37°C, and genotyped by PCR using the kanamycin-cassette-specific primers (forward primer 5’-CGGTGCCCTGAATGAACTGC-3’, reverse primer 5’-CAGTCATAGCCGAATAGCCT-3’) and genomic primers for individual bacterial mutants, which were designed at starting 100-300 bases upstream of start-codon (forward primer) or downstream of stop codon (reverse primer). The correct fragment sizes were verified.

### Quantification of *Pdaf-7::gfp*

*Pdaf-7::gfp* animals at the L4 stage were transferred onto behavioral assay plates seeded with *E. coli* wild-type (BW25113) or test bacteria and incubated at 21°C for 16 hours. Animals mounted on 2% agarose pads and 20 mM sodium azide were imaged under the same parameters (63x objective, Z-stack and exposure time) on the Zeiss Axioimager Z1. Quantification of GFP intensity was performed on maximum fluorescence intensity projections of ASI and ASJ neurons in FIJI.

### Fluorescent reporter quantification assays

All animals containing fluorescent markers (*hsp-6p::gfp*, *hsp-4p::gfp*, *gst-4p::gfp*, *clec-60p::gfp*, *irg-1p::gfp*, or *irg-5p::gfp*) were picked onto behavioral assay plates seeded with *E. coli* test bacteria at the L4 stage and incubated at 21 °C for 16 hours, except animals with the stress reporter *gst-4::gfp*, which were incubated for 8 hours before imaging. Animals mounted on 2% agarose pads and 20 mM sodium azide were imaged on the Zeiss Axioimager Z1. As these reporters are broadly expressed, GFP fluorescence across the whole body was quantified in FIJI.

### Bacterial colonization assay

Animals were treated under behavioral assay condition with indicated bacterial mutants for 12 hours, picked into tubes with M9 buffers, washed three times, and placed on empty NGM plates containing 100μg/mL carbenicillin for 1 hour. At least five animals were then individually picked into a new Eppendorf tube with M9 buffer and homogenized using a motorized tissue grinder (Fisher Scientific) coupled with a 1.5mL plastic pestle (Fisher Scientific). The lysate was plated onto LB plates and incubated at 37°C for 16 hours. Bacterial colonies were counted manually. Three independent biological repeats were performed for *E. coli* wild-type or mutants.

### Pharyngeal pumping assay

Animals were incubated as above for 12 hours on plates seeded with indicated bacteria, then recorded using a LEICA MZ6 microscope coupled with cameras for ten seconds and the number of pharyngeal contractions per animal was counted manually. At least six animals were recorded in each of three independent repeats.

### Brood size and development assay

For the brood size assay, an individual animal at the L4 stage was picked onto one NGM plate seeded with 200 μL bacteria grown under the conditions used in the behavioral assay. Animals were picked onto new plates every day and maintained at 20°C. Brood size was determined by counting the total number of progeny across all plates for each animal. At least three independent biological replicates (6∼8 animals for each repeat) were performed.

For the development assay, five synchronized adult worms were allowed to lay eggs for 3 hours at 20°C, on NGM plates with bacteria grown under the conditions used in the behavioral assay. Eggs were allowed to develop at 20°C for 72 hours, and scored for the number of progeny that had reached the L4 or adult stage as a fraction of all progeny.

### Lifespan assay

Seeded NGM plates used for lifespan assays were prepared with bacteria grown under the conditions used in the behavioral assay. About 20 synchronized L4 worms were picked onto assay plates and maintained at 25°C. To separate parents from offspring, adult animals were picked onto freshly prepared assay plates daily until no further progeny were observed. Dead animals were identified by non-response to gentle prodding. Animals that were lost or crawled off agar were censored.

### Chemical supplementation

50% glucose (Millipore Sigma), 0.2 M cysteine, 0.2 M O-acetyl-serine and 40 mM PLP were prepared by dissolving powders in deionized water, filtered and stored at −20°C. For behavioral analysis with pre-adding chemicals, stock solutions were supplemented into NGM agar immediately before pouring to desired final concentrations (glucose, 0.4%; Cys, 200μM; OAS, 200μM; PLP, 40μM). Bacterial seeding and behavioral assay were performed as described in the section ‘Screen for *C. elegans* aversion to *E. coli* Keio knockout mutants’. For behavioral analysis with post-adding chemicals, assay plates were prepared by first seeding 12.5μL indicated bacterial cultures onto NGM agar for growth, as described in the section ‘Screen for *C. elegans* aversion to *E. coli* Keio knockout mutants’. One hour before the behavioral assay, chemicals were directly added onto the bacterial spots to reach the final concentration under a total volume of 6 mL NGM agar. NGM plates without any added chemicals were assayed in parallel.

### Neuron silencing by histamine assay

1M histamine dihydrochloride (Millipore Sigma) were prepared by dissolving powder in deionized water, filtered and stored at −20°C. Histamine stock solution was then added into NGM agar to a final concentration of 10 mM prior to seeding with bacteria. For behavioral analysis, NGM plates with or without histamine, and animals with or without expression of HisCl1 channels were assayed in parallel.

### Scanning Electron Microscopy

Bacteria were gently eluted from assay plates and fixed in 2% glutaraldehyde, 4% formaldehyde in 0.1 M sodium cacodylate buffer pH 7.2, for more than 1 hour at room temperature followed by overnight fixation at 4 °C. After post-fixation with 1% osmium tetroxide in 0.1M sodium cacodylate buffer, pH 7.2 for 1 hour at room temperature, samples were dehydrated in a graded series of ethanol followed by Hexamethyldisilazane. Bacteria were spread onto glass coverslips and let air-dry overnight. The next morning, samples were sputter-coated with 10 nm of iridium using a Leica ACE600 sputter coater. Images were acquired in a scanning electron microscope JEOL JSM-IT500HR at 5.0 kV (JEOL USA, Inc).

## QUANTIFICATION AND STATISTICAL ANALYSIS

### Statistical analysis

Statistical analyses were performed using GraphPad Prism and are detailed in the figure legends. Detailed statistical summary can be found in *Supplemental Information*.

## ACKNOWLEDGEMENTS

We thank members of the Bargmann lab for their comments on the manuscript. We thank Robert Horvitz, William Schafer, Sreekanth Chalasani, and the Caenorhabditis Genetics Center (P40 OD010440) for sharing strains. We also thank Dr. Hilda Amalia Pasolli at the Rockefeller University Electron Microscopy Facility for conducting electron microscopy. L.F. was supported by Kavli Foundation. C.I.B was supported by a grant from the Chan Zuckerberg Initiative.

## AUTHOR CONTRIBUTIONS

L.F. and C.I.B. designed experiments. L.F. conducted the screen, behavioral, genetic and molecular experiments. J. M-S. and L.Y. conducted behavioral experiments and generated transgenic strains. A.H. generated and validated *tph-1* knockout strains. Y.G. generated NSM::HisCl1 transgenic strains. L.F. and C.I.B. analyzed and interpreted results and wrote the paper with input from all authors.

## INCLUSION AND DIVERSITY

We support diverse, inclusive, and equitable conduct of research.

---

**Figure S1.**
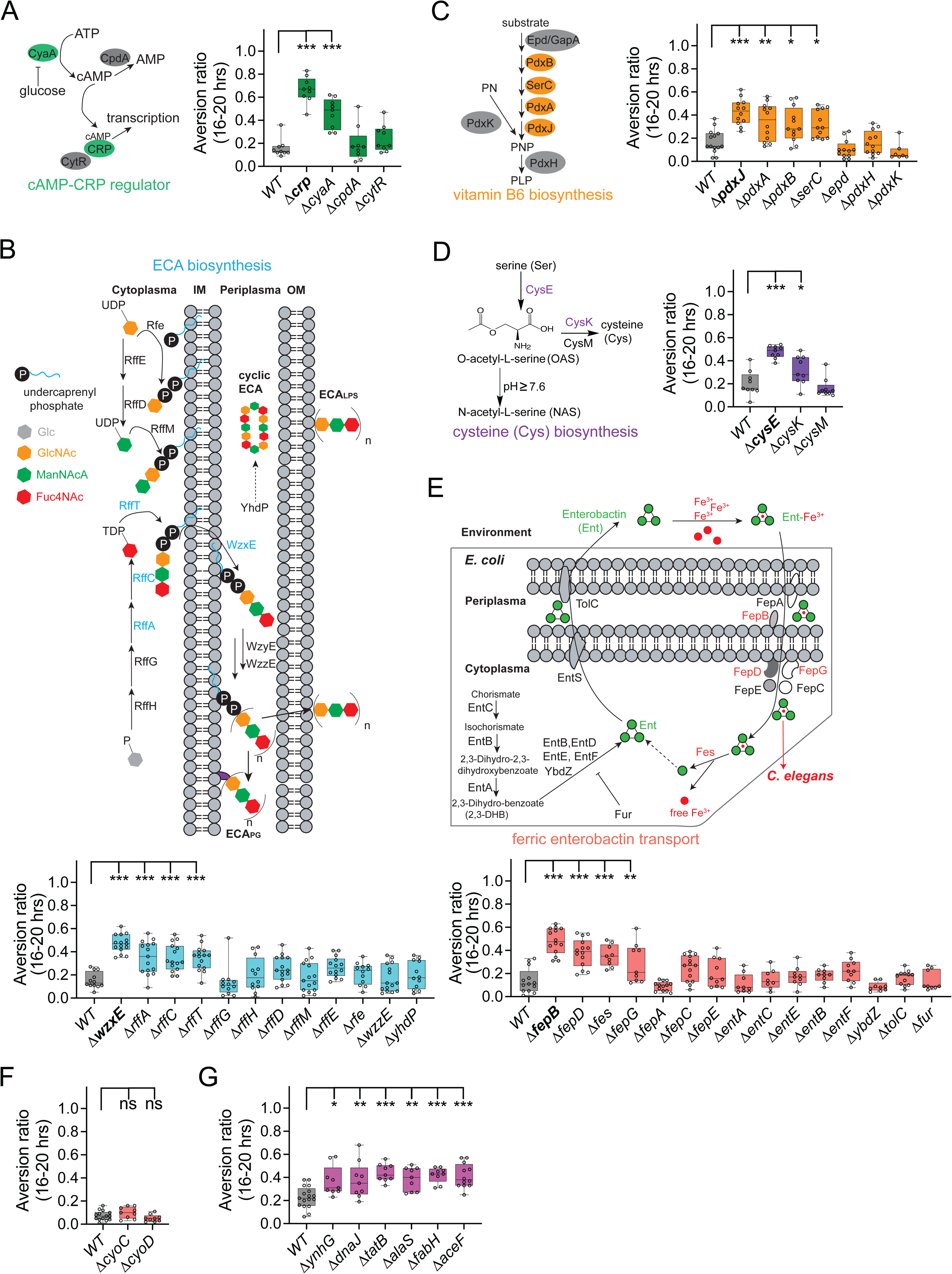
*C. elegans* behavioral responses to *E. coli* mutants identified from the genome-wide screen as well as other genes from the same pathways. Related to Figure 1 and Table 1. (A-E) Complete metabolic pathways and behavioral responses to additional *E. coli* mutants in each pathway, expanded from Figure 1D. (F) Behavioral responses to ROS-producing *E. coli* mutants. (G) Behavioral responses to additional *E. coli* mutants from Table 1. Each data point indicates individual assay. Results are shown with median ± quartiles in boxes and Min to Max whiskers. ns, not significant, *P<0.05, **P<0.01, ***P<0.0001 by One-Way ANOVA, corrected by Dunnett’s multiple comparisons. Statistical analysis with significance is annotated, and for other *E. coli* mutants that are not annotated, not significant.

**Figure S2.**
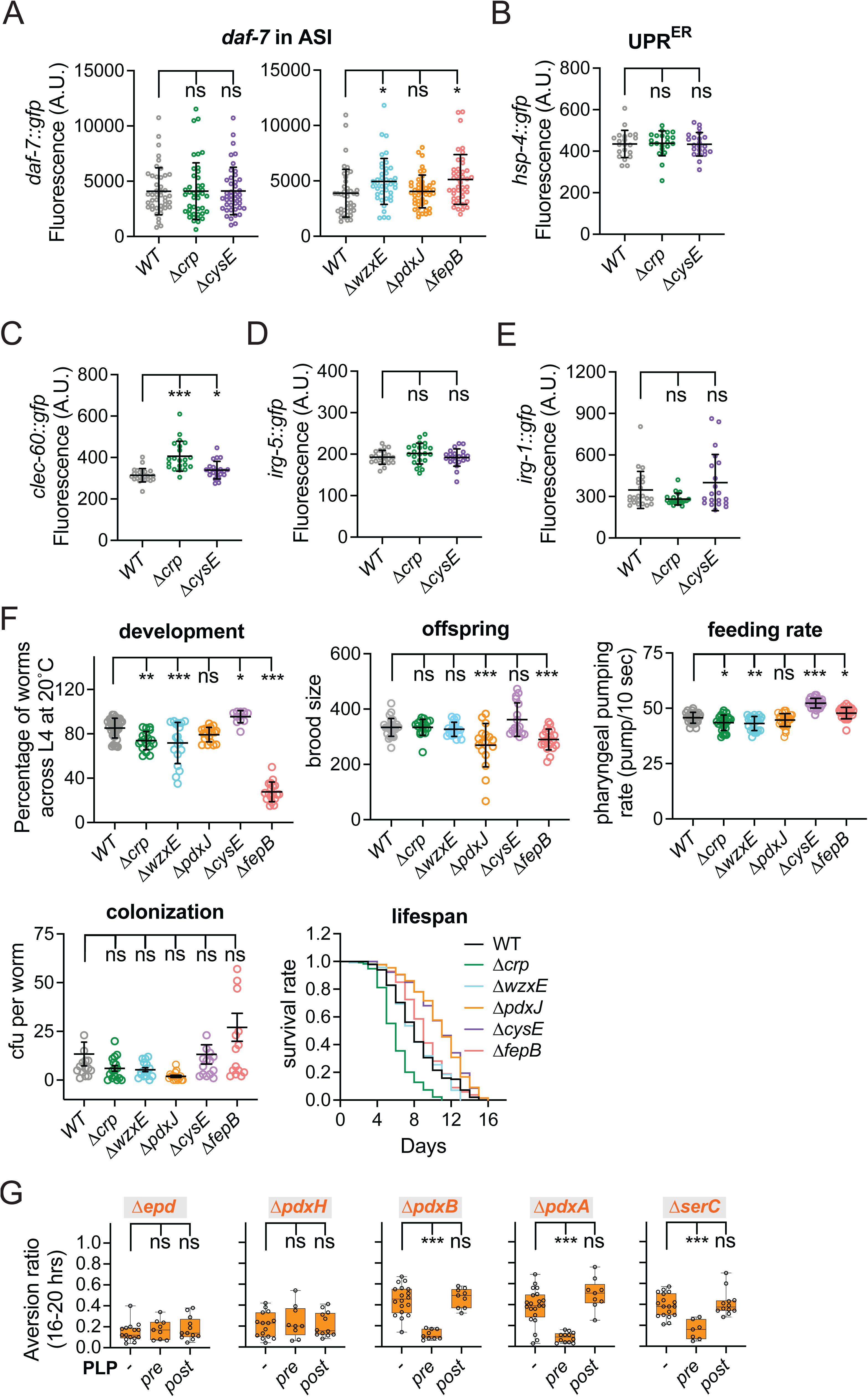
Effects of mediocre diets on animal physiology. Related to Figure 2. (A-E) Effects of bacterial diets on *C. elegans* stress reporters, including *daf-7* expression in ASI (*daf-7::gfp*) (A), UPR^ER^ (*hsp-4::gfp*) (B), and the immune-response genes *clec-60::gfp* (C), *irg-5::gfp* (D), and *irg-1::gfp* (E). GFP fluorescence intensities were quantified using Arbitrary Units (A.U.). Each data point indicates stress response from individual animal. Error bars indicate mean ± SD. ns, not significant, *P<0.05, ***P<0.001 by One-Way ANOVA, corrected by Dunnett’s multiple comparisons. (F) Effects of bacterial diets on *C. elegans* development, brood size, feeding rate, colonization and lifespan. For brood size (offspring), each data point represents the number of progeny from an individual animal. For colonization, each data point represents the number of bacteria isolated from an individual animal. ns, not significant, ***P<0.0001 by One-Way ANOVA, corrected by Dunnett’s multiple comparisons. For feeding rate and development, three biological assays were conducted per condition, including at least five replicates per assay. Each data point indicates individual replicate. Error bars indicate mean ± SD. ns, not significant, *P<0.05, **P<0.01, ***P<0.001 by One-Way ANOVA, corrected by Dunnett’s multiple comparisons. Lifespan assay was performed at 25 °C. (G) Chemical supplementation with PLP. Related to Figure 2F. Only pre-adding PLP suppressed aversion induced by *E. coli* mutants Δ*pdxB*, Δ*pdxA*, or Δ*serC*. 40 μM PLP was used for all experiments. Each data point indicates individual assay. Results are shown with median ± quartiles in boxes and Min to Max whiskers. ns, not significant, ***P<0.0001 by One-Way ANOVA, corrected by Dunnett’s multiple comparisons.

**Figure S3.**
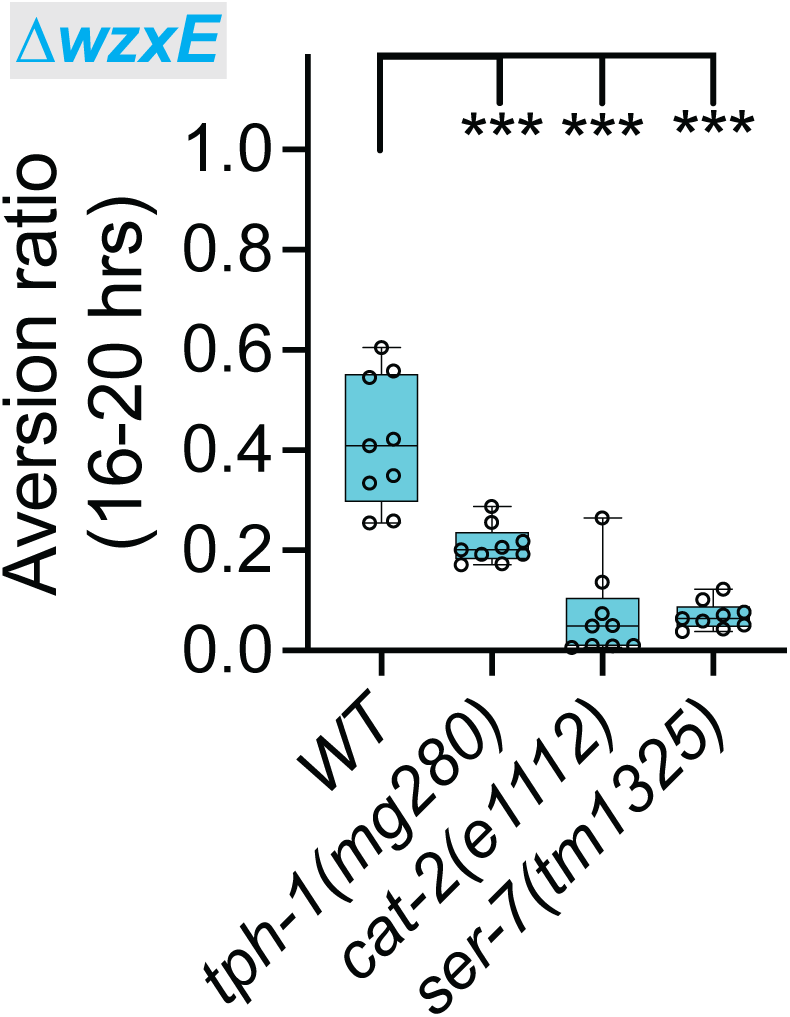
Genetic analysis of *C. elegans* aversion to the mediocre Δ*wzxE* diet. Related to Figure 6. (A) Quantification of aversion ratios of WT, *tph-1*, *cat-2* and *ser-7* animals on the mediocre Δ*wzxE* diet. Each data point indicates individual assay. Results are shown with median ± quartiles in boxes and Min to Max whiskers. ***P<0.001 by One-Way ANOVA, corrected by Dunnett’s multiple comparisons.

**Figure S4.**
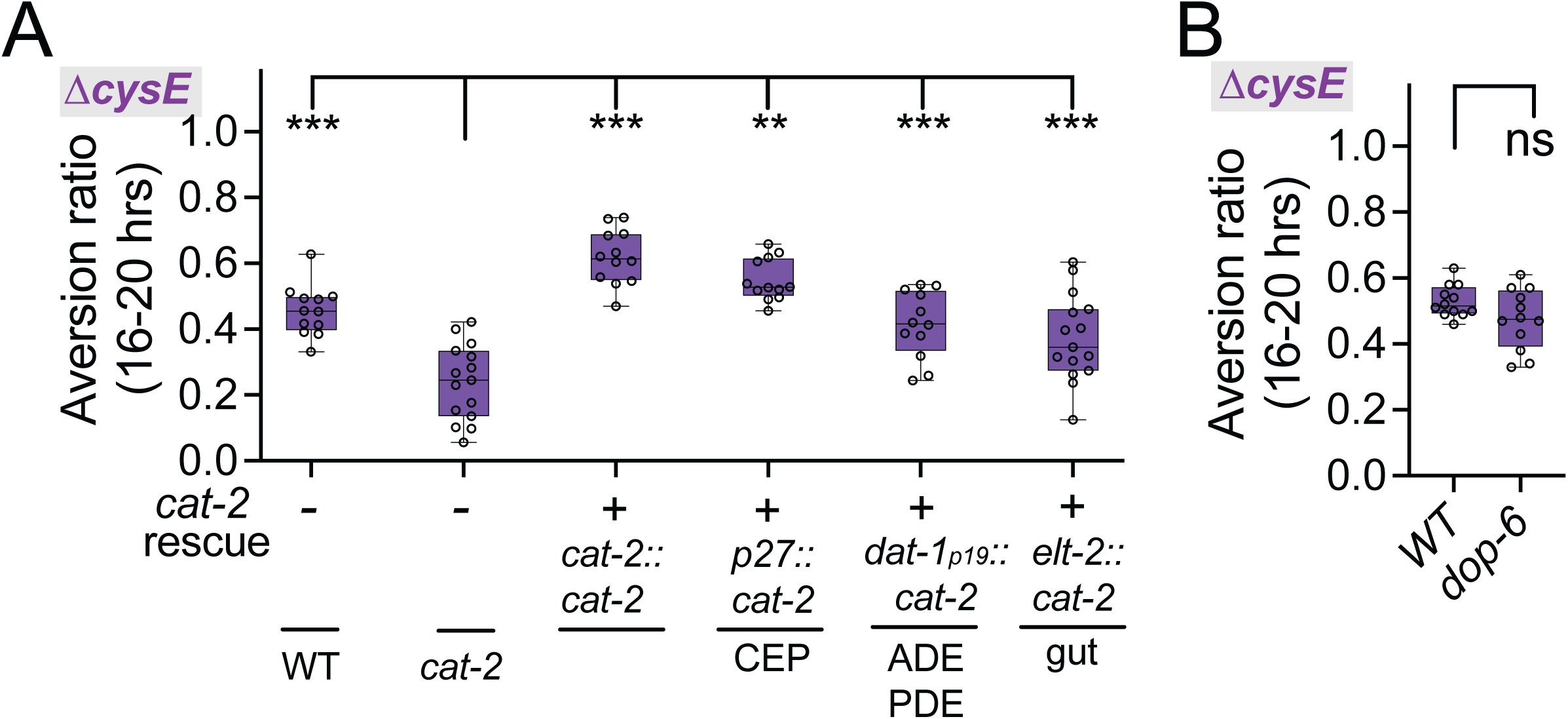
Function of dopaminergic signaling on the mediocre Δ*cysE* diet. Related to Figure 7. Quantification of aversion ratios of *cat-2* rescue lines (A) or *dop-6* mutant (B) on the Δ*cysE* mediocre diet. Each data point indicates individual assay. Results are shown with median ± quartiles in boxes and Min to Max whiskers. ns, not significant, **P<0.01, ***P<0.001 by One-Way ANOVA, corrected by Dunnett’s multiple comparisons (panel A) and two-tailed, unpaired t test (panel B, *dop-6*).

